# MiniBAR/KIAA0355 is a dual Rac and Rab effector required for ciliogenesis

**DOI:** 10.1101/2023.07.24.550339

**Authors:** Ronan Shaughnessy, Murielle Serres, Sophie Escot, Hussein Hammich, Frédérique Cuvelier, Audrey Salles, Murielle Rocancourt, Quentin Verdon, Anne-Lise Gaffuri, Yannick Sourigues, Gilles Malherbe, Leonid Velikovsky, Florian Chardon, Jean-Yves Tinevez, Isabelle Callebaut, Etienne Formstecher, Anne Houdusse, Nicolas David, Olena Pylypenko, Arnaud Echard

**Affiliations:** Institut Pasteur, Université de Paris, CNRS UMR3691, Membrane Traffic and Cell Division Lab, 25-28 rue du Dr Roux, F-75015 Paris, France; Laboratoire d’Optique et Biosciences (LOB), CNRS, INSERM, Ecole Polytechnique, Institut Polytechnique de Paris, 91120 Palaiseau, France; Institut Curie, PSL Research University, CNRS UMR144, Structural Motility, 26 rue d’Ulm, F-75005 Paris, France; Institut Pasteur, Université de Paris, UTechS Photonic BioImaging (UTechS PBI), Centre de Recherche et de Ressources Technologiques C2RT, 25-28 rue du Dr Roux, F-75015 Paris, France; Institut Pasteur, Université de Paris, Image Analysis Hub, 25-28 rue du Dr Roux, F-75015 Paris, France; Sorbonne Université, Muséum National d’Histoire Naturelle, UMR CNRS 7590, Institut de Minéralogie, de Physique des Matériaux et de Cosmochimie, IMPMC, Paris, France; Hybrigenics Services SAS, 1 rue Pierre Fontaine 91000 Evry – Courcouronnes, France

**Keywords:** Rab GTPase, Rac GTPase, Rab35, BAR domain, membrane traf-ficking, actin cytoskeleton, Myosin II, ciliogenesis, ciliopathies

## Abstract

Cilia protrude from the cell surface and play critical roles in intracellular signaling, environmental sensing and development. Actin-dependent contractility and intracellular trafficking are both required for ciliogenesis, but little is known about how these processes are coordinated. Here, we identified a Rac1-and Rab35-binding protein with a truncated BAR domain that we named MiniBAR (aka KIAA0355/GARRE) which plays a key role in ciliogenesis. MiniBAR colocalizes with Rac1 and Rab35 at the plasma membrane and on intracellular vesicles trafficking to the ciliary base and exhibits remarkable fast pulses at the ciliary membrane. MiniBAR depletion leads to short cilia resulting from abnormal Rac-GTP/Rho-GTP levels, increased acto-myosin-II-dependent contractility together with defective trafficking of IFT88 and ARL13B into cilia. MiniBAR-depleted zebrafish embryos display dysfunctional short cilia and hallmarks of ciliopathies including left-right asymmetry defects. Thus, MiniBAR is a unique dual Rac and Rab effector that controls both actin cytoskeleton and membrane trafficking for cili-ogenesis.

## Introduction

Flagella and cilia are evolutionarily conserved microtubule-based protrusions essential for cell locomotion, fluid movement and sensing of extracellular cues (Anvarian et al., 2019; Goetz and Anderson, 2010; Wheway et al., 2018). Primary cilia are found at the surface of most vertebrate cells and function as cellular antennas that detect various chemical and mechanical signals, thereby playing crucial roles in intracellular signaling and development (Anvarian et al., 2019; Goetz and Anderson, 2010; Wheway et al., 2018). Motile cilia in the embryonic node control left-right patterning of the body plan, in particular by generating a directional fluid flow essential for symmetry breaking of key transcription factors (Hirokawa et al., 2006). Strikingly, dozens of proteins are required for the formation and maintenance of cilia (Kim et al., 2010; Wheway et al., 2015). Mutations of the corresponding genes in humans can lead to severe diseases known as cil-iopathies in up to 1 out of 725 individuals (Smith et al., 2020; Wheway et al., 2019), which can cause various pathologies including polycystic kidney diseases, nephrophtisis, hetero-taxis and situs inversus, and Meckel-Gruber, Bardet-Biedl or Joubert syndromes (Hildebrandt et al., 2011; Mitchison and Valente, 2017; Reiter and Leroux, 2017). Primary cilia consist of a membrane evagination protruding from the cell surface that surrounds a microtubule-based axoneme nucleated by a basal body (Malicki and Johnson, 2017; Satir et al., 2010). Depending on the cell type, the basal body either docks to the plasma membrane and the primary cilium directly growths from this location, or the axoneme starts to grow into an intracellular ciliary vesicle which subsequently fuses to the plasma membrane, as in RPE-1 (retinal pigment epithelial) cells (Malicki and Johnson, 2017; Molla-Herman et al., 2010; Sorokin, 1968). In both pathways, the assembly, growth, maintenance and proper functioning of the cilium requires the selective delivery of membrane and cytosolic cargoes within the cilium. This relies on both highly conserved intraflagellar transport (IFT) and intracellular membrane trafficking (Klena and Pigino, 2022; Mul et al., 2022). Accordingly, Rab GTPases, which are key regulators of membrane trafficking in eukaryotic cells (Homma et al., 2021), play a pivotal role in cilium biology, with at least ten Rab proteins involved in cilium formation, function and composition (Blacque et al., 2018; Madhivanan and Aguilar, 2014). For instance, a Rab11-Rabin8-Rab8 cascade and their effectors (EHD1, MICAL-L1 and the exocyst), together with the coat and adaptor complex BBSome, control trafficking to the ciliary vesicle and cilium growth (Feng et al., 2012; Knödler et al., 2010; Lu et al., 2015b; Nachury et al., 2007; Westlake et al., 2011; Wingfield et al., 2018; Xie et al., 2019). In parallel, the actin cytoskeleton and actin binding proteins participate in several aspects of cilia formation, elongation and shedding (Hoffman and Prekeris, 2022; Magistrati et al., 2022; Nager et al., 2017; Ojeda Naharros and Nachury, 2022; Phua et al., 2017; Smith et al., 2020; Wu et al., 2018). Importantly, excessive actin polymerization and contractility impair ciliogenesis possibly by imposing a physical barrier to cilia-targeted vesicle transport and by increasing cortical or membrane tension that prevents outward growth of cilia (Hoffman and Prekeris, 2022; Smith et al., 2020). Consequently, pharmacological inhibition of F-actin polymerization and of acto-myosin II-dependent contractility can enhance ciliogenesis in normal cells or restore normal ciliogenesis in cells depleted of important ciliary proteins (Kim et al., 2015; Kim et al., 2010; Pitaval et al., 2010). Thus, both intracellular trafficking and the actin cytoskeleton dynamics play critical roles in cilia initiation and growth. Yet, little is known about how these two important cellular processes are coordinated during ciliogenesis. Rab35 is a plasma membrane and endosomal Rab GTPase implicated in diverse cellular functions, ranging from cytokinesis, phagocytosis, cell migration and neurite outgrowth to autophagy (Chaineau et al., 2013; Klinkert and Echard, 2016; Shaughnessy and Echard, 2018). Recently, Rab35 was reported to control cilia length and function, both in human cultured cells and in zebrafish embryos, but the Rab35 effectors involved remain unknown (Kuhns et al., 2019). Here, we identified MiniBAR as a unique Rab35 and Rac1 effector with a truncated BAR domain that plays a critical role in ciliogenesis in RPE-1 cells and zebrafish embryos. We demonstrate that the interaction of MiniBAR with each GTPase is necessary for proper cilia growth. We also show that this dual Rab/Rac effector promotes ciliogenesis by controlling and coordinating both membrane trafficking of key ciliary cargoes and acto-myosin II-dependent cellular contractility.

## Results

### MiniBAR is a dual Rac1 and Rab35 effector with a truncated BAR domain

Using both GTP-locked mutants of Rab35 (Rab35Q67L) and of Rac1 (Rac1G12V) as baits in yeast two hybrid screens, we identified the poorly characterized human protein KIAA0355 among the top hits (see Methods), indicating that KIAA0355 might be a dual Rab35 and Rac1 effector protein. Interestingly, while we were studying KIAA0355, this protein was found through systematic Rho-family GTPase proximity interaction assays and named GARRE for "Granule-associated Rac and RhoG effector protein", since it directly interacted in vitro with Rac1G12V and potentially with other Rac-family members -Rac2, Rac3 and RhoG (Bagci et al., 2020). However, neither the function of this 1070 aa protein conserved in Vertebrates (**Figure 1A and S1A**), nor the relevance of its interaction with Rac1 were known. Based on its structure and localization (see below), we propose to name KIAA0355/GARRE as MiniBAR, and we will use this alias throughout this study.

**Fig. 1.**
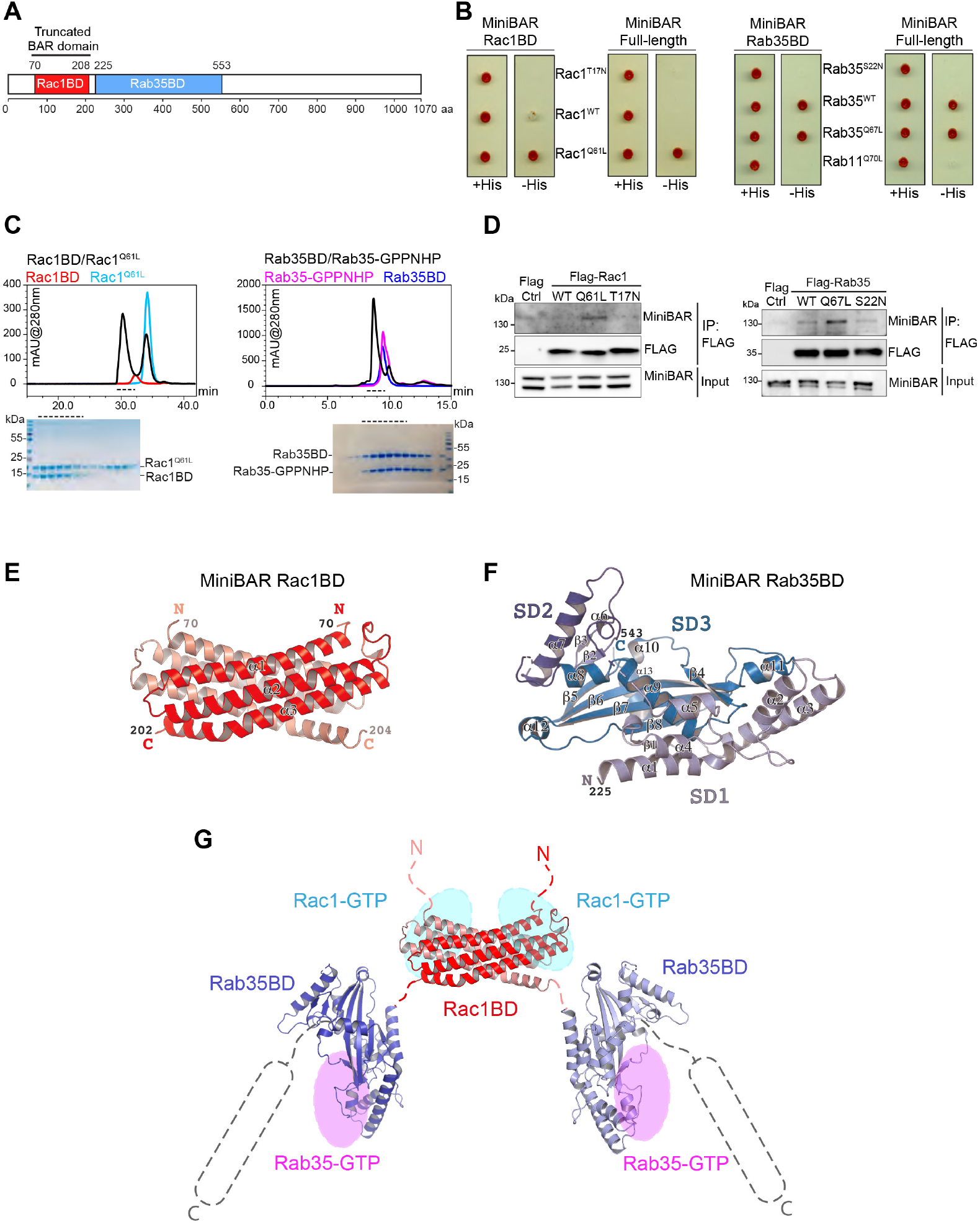
MiniBAR is a dual Rac1 and Rab35 effector with a truncated BAR domain. (**A**) Scheme representing MiniBAR domains. (**B**) S. cerevisiae L40 reporter strain was transformed with plasmids encoding Gal4 Activation Domain (GAD) fused to MiniBAR to examine interactions with LexA fused to indicated Rac1 or Rab35 mutants. Growth on a medium without histidine (-His) indicates an interaction with the corresponding proteins. (**C**) Analytical gel-filtration elution profiles of Rac1BD (left panels), Rab35 BD (right panels), Rab35-GPPNHP and Rac1Q61L in comparison with their complexes. The elution fractions corresponding to the complexes elution peaks (black dashed lines) were analyzed by SDS-PAGE, and masses were determined by MALS. Rac1BD (MassMALS 30.1 ± 0.4 percent kDa), Rac1Q61L (MassMALS 20.6 ± 0.8 percent kDa), Rac1BD/Rac1Q61L complex (MassMALS 69.9 ± 0 .3 percent kDa), Rab35BD (MassMALS 36.4 ± 0.7 percent kDa), Rab35-GPPNHP (MassMALS 22.2 ± 0.1 percent kDa), Rab35BD/Rab35-GPPNHP complex (MassMALS 50.8 ± 0.8 percent kDa). (**D**) Flag Immunoprecipitation (IP Flag) from RPE-1 cells transfected with plasmids encoding Flag alone, Flag-tagged Rac1 or Rab35 constructs. Flag-proteins and co-immunoprecipitated endogenous MiniBAR were detected by western blot using anti-MiniBAR antibodies and anti-Flag antibodies. Bottom panels: inputs (3 of total lysates). (**E**) Crystal structure of Rac1BD: 3 antiparallel helix unit forms a homodimer in an antiparallel fashion resulting in a 6-helix bundle. (**F**) Crystal structure of Rab35BD composed of 3 subdomains (SD-1,-2,-3). (**G**) Model of MiniBAR Rac1BD-Rab35BD unit, the putative positions of Rac1, Rab35 binding sites and the C-terminal disordered domain are indicated (see also Figure S5).

We first delimited the Rac1 binding domain or Rac1BD (aa 70-208, MiniBAR70-208) and the Rab35 binding domain or Rab35BD (aa 225-553, MiniBAR225-553) using yeast two hybrid assays (**Figure 1A-B**). Each domain as well as the full-length protein interacted selectively with Rac1 and Rab35 mutants locked in their GTP-bound states, but not with mutants locked in their GDP-bound states (**Figure 1B**). Among the 55 Rab GTPases tested, only Rab35 interacted with MiniBAR, suggesting that MiniBAR is a Rab35-specific effector (**Figure S1B**). Gel filtration experiments with recombinant, purified proteins demonstrated that Rac1BD and Rab35BD assembled into complexes with the active forms of Rac1 and Rab35, respectively (**Figure 1C**). Importantly, Rac1 and Rab35 also interacted individually and simultaneously with MiniBAR70-553 containing the two binding domains, showing that MiniBAR is a dual Rac-Rab effector (**Figure S1C-D**). Isothermal titration calorimetry (ITC) revealed that MiniBAR interacted with GTP-bound -but not GDP-bound-Rac1 and Rab35, with Kds of approximately 0.2 µM and 2 µM, respectively (**Figure S1E**). Finally, preferential binding of endogenous MiniBAR to GTP-bound Rac1 and Rab35 was validated by coimmunoprecipitation experiments from RPE-1 cells (**Figure 1D**). Altogether, MiniBAR interacts selectively with GTP-bound Rac1 and Rab35 via two distinct, adjacent domains.

The Rac1BD overlaps with a domain of unknown function 4745 (DUF4745) in the Pfam database that is predicted to form a BAR (Bin/amphiphysin/Rvs) domain, which is known to sense membrane curvature (Bagci et al., 2020; Carman and Dominguez, 2018; Simunovic et al., 2015) (**Figure S2A**). Intriguingly, we noticed that the helices that form "extended arms" in the canonical BAR-domain of Amphiphysin (**grey in Figure S2A**) were shorter in DUF4745 and MiniBAR (**red in Figure S2A**). Accordingly, the crystal structure of the Rac1BD that we solved (**Figure 1E**, **Movie 1 and Table S1**) revealed an unusual, truncated or "Mini" BAR domain consisting of a 3-helix-based homodimer, with strong similarity to the central part of a canonical BAR-domain (Carman and Dominguez, 2018; Simunovic et al., 2015) (**Figure S2B, top panels**). The "Mini" BAR domain surface has a positively charged cluster that might mediate interactions with negatively charged membranes (**Figure S2B, lower panels**). However, this cluster is condensed in the middle part of the MiniBAR homodimer, which differs from the more peripheral distribution in canonical BAR domains (**Figure S2B**). Small-Angle X-ray Scattering (SAXS) experiments of the Rac1BD are consistent with the formation of a homodimer in solution (**Figure S2C and Table S1**). In addition, the Rac1BD : Rac1 complex molecular mass defined by MALS corresponds to 2 Rac1 : 1 Rac1BD homodimer stoichiometry (**Table S1**).

The crystal structure of the Rab35BD that we determined revealed that it comprises three subdomains (SD-1,2,3), with the topologically central subdomain SD-3 exhibiting a Cystatin/Monellin characteristic protein fold (**Figure 1F, S3A-C, Movie 1 and Table S1**). SAXS further showed that the Rab35BD is monomeric in solution, with the last 40 residues (including helix alpha 13 and a preceding loop) being flexible (**Figure S2D and Table S1**). Furthermore, oligomerization of the Rac1BD resulted in the dimerization of the entire Rac1BD-Rab35BD unit (MiniBAR70-553) (**Table S1**). Finally, SAXS suggested that different orientations of two Rab35BDs with respect to the central Rac1BD dimer are likely present in solution (**Figure S2E**). This might enable interactions with membrane-bound Rac1 and Rab35 GTPases present either on the same membrane (in cis) or on two differ-ent/opposite membranes (in trans). Based on these biochem-ical and structural evidence, we conclude that MiniBAR is a Rac1 and Rab35 binding protein and propose a structural model for the overall organization of the complex (**Figure 1G and results below for Rac/Rab interfaces**). To our knowl-edge, MiniBAR represents the only known dual Rho-family and Rab-family interacting protein with a truncated BAR do-main.

### MiniBAR partially colocalizes with Rac1 and Rab35 at the plasma membrane and on dynamic intracellular vesi-cles

Rac1 localizes at the plasma membrane and on intra-cellular compartments, including endosomes, where it ac-tivates actin polymerization (Gautreau et al., 2022; Phuyal and Farhan, 2019). In conditions that do not induce cilia formation, time-lapse spinning-disk confocal microscopy re-vealed that fluorescently tagged MiniBAR (stably expressed MiniBAR-GFP) largely colocalized with Rac1 at plasma membrane ruffles in RPE-1 cells (**Figure 2A and Movie 2**). MiniBAR also partially colocalized with Rac1 on dynamic intracellular vesicles, with approximately 60 percent of Mini-BAR vesicles being positive for Rac1 (**Figure 2A and Movie 2**).

**Fig. 2.**
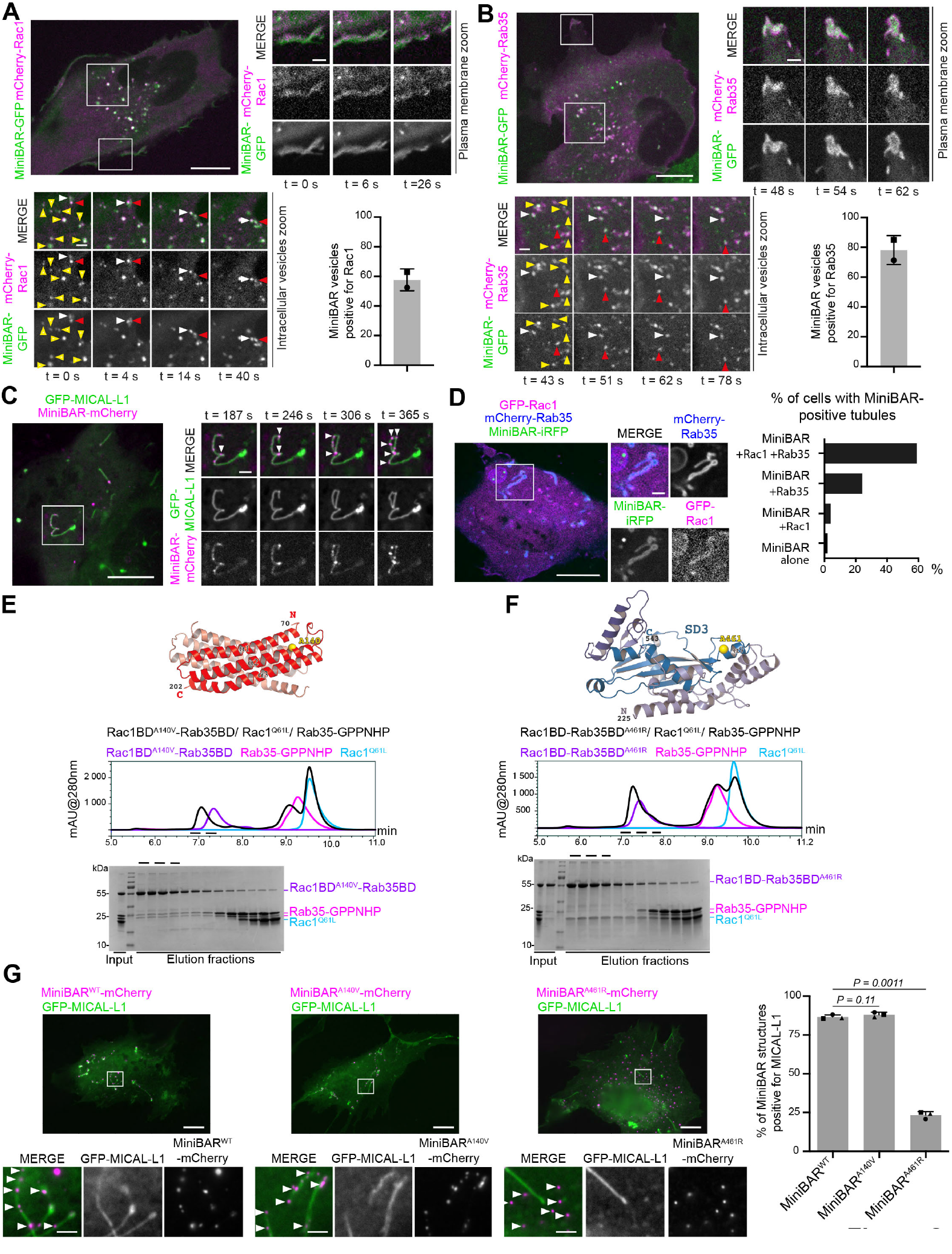
MiniBAR partially colocalizes with Rac1 and Rab35 at the plasma membrane and on dynamic intracellular vesicles. (**A**) Top left panel: Snapshot of a movie of RPE-1 cells stably co-expressing MiniBAR-GFP and mCherry-Rac1. Top right and Bottom left panels: Selected time points of zoomed regions showing the plasma membrane and intracellular vesicles, respectively. Arrowheads: co-localization on vesicles; yellow and red arrowheads: displacement of two selected moving vesicles. Scale bars, 10 µm (general view) and 2 µm (insets). Bottom right panel: percentage of MiniBAR-GFP vesicles positive for mCherry-Rac1. Mean ± SD, n = 57-124 MiniBAR-positive vesicles per cell, n = 14 cells.(**B**) Same as (A) for an RPE-1 cells co-expressing MiniBAR-GFP and mCherry-Rab35. Scale bars, 10 µm (general view) and 2 µm (insets). Bottom right panel: percentage of MiniBAR-GFP vesicles positive for mCherry-Rab35. Mean ± SD, n = 91-143 MiniBAR-positive vesicles per cell, n = 6 cells.(**C**) Left panel: Snapshot of a movie of an RPE-1 cells co-expressing GFP-MICAL-L1 and MiniBAR-mCherry. Right panels: snapshots of a zoomed region. Arrowheads: accumulation of MiniBAR. Scale bars, 10 µm (general view) and 2 µm (insets).(**D**) Left panels: RPE-1 cell co-expressing GFP-Rac1, mCherry-Rab35 and MiniBAR-iRFP. Zoomed region: tubular structure labelled by the three proteins. Scale bars, 10 µm (general view) and 2 µm (insets). Right panel: percentage of cells with MiniBAR-GFP-positive tubules after expression or not of Rab35 and/or Rac1, as indicated. n > 50 cells per condition.(**E-F**) Analytical gel-filtration elution profiles of Rac1BD-Rab35BD (MiniBAR70-553) mutants, Rab35-GPPNHP and Rac1Q61L in comparison with the ternary protein complexes. The elution fractions corresponding to the complex elution peaks were analyzed by SDS-PAGE. (**E**) Yellow ball: A140V mutation mapped on the Rac1BD structure. The Rac1BDA140V-Rab35BD mutant co-elutes with Rab35 but not with Rac1. (**F**) Yellow ball: A461R mutation mapped on the Rab35BD structure. The Rac1BD-Rab35BDA461R mutant co-elutes with Rac1 but not with Rab35.(**G**) Left panels: fixed RPE-1 cells co-expressing GFP-MICAL-L1 with either MiniBARWT-mCherry, MiniBARA140V-mCherry or MiniBARA461R-mCherry, as indicated. Scale bars, 10 µm (general view) and 2 µm (insets). Right panel: percentage of MiniBAR-GFP structures positive for MICAL-L1 in the indicated conditions. Mean ± SD, n = 61-96 MiniBAR-positive vesicles per cell, n = 33-36 cells per condition, N = 3 independent experiments. Paired student t-test.

Rab35 is known to localize at the plasma membrane and on a particular class of intracellular recycling endosomes (Allaire et al., 2010; Allaire et al., 2013; Cauvin et al., 2016; Kobayashi et al., 2014b; Kobayashi and Fukuda, 2013; Kouranti et al., 2006; Rahajeng et al., 2012). As observed with Rac1, MiniBAR-GFP colocalized with Rab35 at the plasma membrane ruffles and protrusions (Figure 2B and Movie 3) and was found on dynamic vesicles that were largely positive (> 75 percent) for Rab35 (**Figure 2B and Movie 3**). MiniBAR/KIAA0355 was recently shown to establish proximity labeling with components (GW182 and Ago2) of membrane-less, phase-separated RNA-containing P-granules (Bagci et al., 2020; Youn et al., 2018). However, co-localization of MiniBAR with P-granules has not been directly investigated. In our experimental conditions and cells, we found that endogenous MiniBAR is present on dynamic membrane vesicles rather than on membrane-less P-granules (**Figure S4A-F and details within**). Consistent with membrane association, MiniBAR-GFP was enriched at the extremities and/or in discrete spots along intracellular tubulo-vesicles labelled by MICAL-L1, a marker of the Rab35 recycling endosomes (Kobayashi et al., 2014b; Rahajeng et al., 2012) (**Figure S4G**). Time-lapse confocal spinning-disk microscopy confirmed the uneven localization of MiniBAR along MICAL-L1 tubules in cells expressing both MICAL-L1-GFP and MiniBAR-mCherry (**Figure 2C and Movie 4**). MiniBAR was enriched where MICAL-L1 tubules experienced strong local deformations (**Figure 2C, arrowheads**), suggesting preferential association of MiniBAR in tubular regions with changes in curvature, perhaps through its unusual BAR domain. Finally, triple co-localization of GFP-Rac1, mCherry-Rab35 and MiniBAR-iRFP was observed on intracellular vesicular structures and at the plasma membrane (**Figure 2D**). Noteworthy, the formation of prominent MiniBAR-positive tubules was induced by the expression of Rab35 and even more potently by its co-expression with Rac1 (**Figure 2D**), suggesting that the three proteins act together to sculpt internal membranes. We conclude that MiniBAR partially colocalizes with Rac1 and Rab35 at the plasma membrane and on dynamic intracellular vesicles in RPE-1 cells.

### Direct MiniBAR / Rab35 interaction is required for vesicular localization of MiniBAR

Guided by our crystal structures (**Figure 1E-F**), we next designed specific point mutations in MiniBAR that selectively disrupted its binding to either Rac1 or Rab35 (**Figure S5A)**. Point mutations A140V (as well as A140T and M144K) in helix alpha 2 of Rac1BD and A461R in helix alpha 11 of Rab35BD abolished the interaction of full-length MiniBAR with Rac1 and Rab35, respectively (**Figure S5A**). The corresponding mutations in recombinant MiniBAR 70-553 (Rac1BD-Rab35BD) also abolished the direct binding to the corresponding GTPase but left unchanged the binding to the other partner, as demonstrated by gel filtration (**Figure 2E-F**). The results of the experimental mutational analysis are consistent with the AlphaFold (Varadi et al., 2022) calculated models of the GTPases/MiniBAR complexes (**Figure S5B**). Thus, A140V in MiniBAR selectively disrupts the interaction with Rac1, while A461R selectively disrupts the interaction with Rab35. Full-length MiniBARA140V-GFP localized to the plasma membrane and remained associated with MICAL-L1 intracellular structures (**Figure 2G**). In contrast, MiniBARA461R-GFP largely lost its association with MICAL-L1 vesicles, while preserving its association with the plasma membrane (**Figure 2G**). We conclude that the interaction of MiniBAR with intracellular vesicles critically depends on its ability to interact with Rab35 but not with Rac1.

### MiniBAR accumulates at the ciliary base and rapidly pulses at the ciliary membrane

Since intracellular trafficking and notably Rab35 have been recently implicated in cilium biology, we next examined MiniBAR localization during ciliogenesis. Immunofluorescence in RPE-1 cells after serum starvation —which induces cilia formation— revealed that endogenous MiniBAR accumulated as vesicular structures surrounding the ciliary base in 100 percent of ciliated cells (n= 157 cells), as shown by super-resolution structured illumination microscopy (**Figure 3A and S6A for staining specificity**). Time course after serum removal showed that MiniBAR staining was present before cilia start to elongate, culminated in the early times of ciliogenesis (2-4 h) but remained at the ciliary base at later steps of ciliogenesis (**Figure 3B**). Videomicroscopy confirmed that trafficking at the ciliary base of MiniBAR-GFP vesicles positive for Rac1 and Rab35 occurred before (**Figure 3C-D**) and during cilia elongation (**Figure 3E-F**). Together, the timing and localization data suggest that MiniBAR vesicles could deliver cargoes in the ciliary base region, before and during cilia elongation. Beside dynamic tubulo-vesicles at the ciliary base (**Figure 3G, yellow arrowhead**), MiniBAR-GFP was usually present at low levels at the plasma membrane, in par-ticular at the ciliary membrane. However, while acquiring movies at 2 second frequencies, we observed striking, tran-sient accumulations of MiniBAR-GFP along cilia, as shown in Figure 3G (white brackets) and Movie 5. MiniBAR-GFP could fill the entire axoneme within 2-4 seconds and accumu-lated strongly in cilia in the next 15 seconds before rapidly disappearing (**Figure 3G**). In this example, 5 distinct Mini-BAR pulses were observed during a 6-min long movie. We could distinctly observe MiniBAR-GFP as lines parallel to the axoneme signal at the resolution of confocal microscopy (**Figure 3G, time point 189 s**), suggesting that MiniBAR-GFP associated with the ciliary membrane. At least one MiniBAR pulse was observed during 6-min long movies in approximately 60 percent of the ciliated cells with clear SiR-Tubulin-positive axonema (**as in Figure 3G**) and analyzed 4- 8 hours after serum removal (**Figure 3H**). Although irregular, one pulse every 1.6 ± 1.16 min (mean ± SD, n= 27 cilia) was detected in pulsating cells in average. When movies were extended to 12 min, all ciliated cells showed at least one MiniBAR pulse along cilia 4-8 hours after serum removal (n = 10/10 cells). In contrast, MiniBAR pulses were ab-sent during 6-min long movies in most ciliated cells recorded 24 hours after serum removal (n= 13/16 cells). Thus, Mini-BAR pulses at the ciliary membrane in all ciliated cells dur-ing early but not late phases of ciliogenesis. mCherry-Rac1 was found at constant levels along the cilium (**Figure 3I**) and MiniBARA140V pulsed in as many cells as wild type Mini-BAR (**Figure 3H**), indicating that Rac1/MiniBAR interaction is not required for MiniBAR pulsatile behavior. In contrast, several lines of evidence indicated that Rab35 drives Mini-BAR pulsation in cilia. First, mCherryRab35 also showed pulsatile behavior, and MiniBAR pulses occurred simulta-neously (**Figure 3j, white brackets, and Movie 6**). Sec-ond, MiniBARA461R-GFP never showed pulsatile behav-ior (**Figure 3H, n= 0/23 cells**). Third, Rab35 depletion es-sentially abolished MiniBAR pulses in cilia (**Figure 3H, n= 1/11 cilia**). Altogether, high-frequency acquisition movies revealed the existence of fast (38 s ± 23.6 s, mean ± SD, n= 35 pulses from 14 cells) pulses of MiniBAR at the cil-iary membrane that critically depend on its interaction with Rab35.

**Fig. 3.**
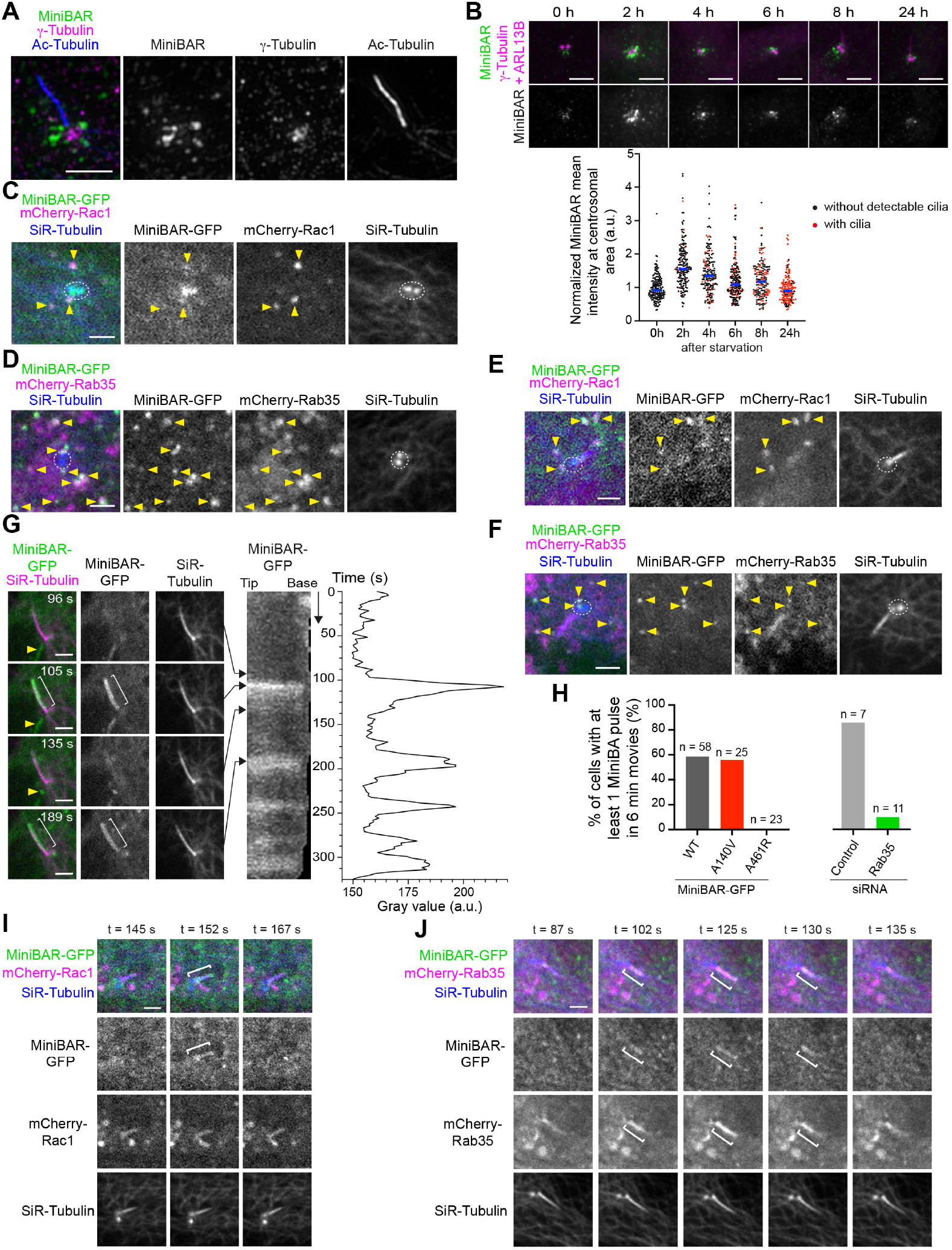
MiniBAR accumulates at the ciliary base and rapidly pulses at the ciliary membrane. (**A**) Structured Illumination Microscopy images of a ciliated RPE-1 cell stained with endogenous MiniBAR, acetylated-Tubulin (to visualize the axoneme) and G-Tubulin (to visualize the basal body). Scale bar, 2 µm.(**B**) Top panels: Endogenous MiniBAR localization in fixed RPE-1 cells following serum removal (time course indicated in hours). Cells were also stained for ARL13B and G-Tubulin to visualize cilia and basal bodies, respectively. Scale bar, 4 µm. Bottom panel: Quantification of MiniBAR intensity at the centrosomal area following serum removal. n = 164-177 cells per condition. N = 3 independent experiments. (**C**) Snapshot of a movie of RPE-1 cells stably co-expressing MiniBAR-GFP and mCherry-Rac1, incubated with SiR-Tubulin (to visualize the microtubules, with a strong accumulation at both centrioles, encircled). Acquisition started 3-4 h after serum removal. Arrowheads: co-localization on vesicles at the centrosome area. Scale bar, 2 µm.(**D**) Snapshot of a movie of RPE-1 cells stably expressing MiniBAR-GFP, transfected with mCherry-Rab35 and incubated with SiR-Tubulin. Acquisition started 3-4 h after serum removal. Arrowheads: co-localization on vesicles at the centrosome area. Scale bar, 2 µm.(**E**) and (**F**) Same as in (C) and (D), respectively, in RPE-1 cells with a detectable cilia. Scale bars, 2 µm.(**G**) Left panels: Snapshots of a movie of RPE-1 cells stably expressing MiniBAR-GFP and incubated with SiR-Tubulin. Acquisition started 3-4 h after serum removal. Yellow arrowhead: a MiniBAR-positive tubulo-vesicle at the ciliary base. White brackets: pulsatile accumulation of MiniBAR-GFP along the cilium. Scale bar, 2 µm. Right panels: Corresponding kymograph of MiniBAR-GFP intensity along the cilium and quantification of the intensity during a 6-min movie, showing 5 pulses of MiniBAR.(**H**) Percentage of cells with at least 1 pulse of MiniBAR-GFP in 6-min movies in RPE-1 cells stably expressing either MiniBARWT-GFP, MiniBARA140V-GFP or MiniBARA461R-GFP, and in RPE-1 cells stably expressing MiniBARWT transfected with either control or Rab35 siRNAs, as indicated. (**I**) Snapshots of a movie of RPE-1 cells stably expressing MiniBAR-GFP, transfected with mCherry-Rac1 and incubated with SiR-Tubulin. Acquisition started 3-4 h after serum removal. White brackets: MiniBAR-GFP pulse along the cilium, while mCherry-Rac1 levels were unchanged. Scale bar, 2 µm.(**J**) Snapshots of a movie of RPE-1 cells stably expressing MiniBAR-GFP, transfected with mCherry-Rab35 and incubated with SiR-Tubulin. Acquisition started 3-4 h after serum removal. White brackets: simultaneous pulsatile accumulation of mCherry-Rab35 and MiniBAR-GFP along the cilium. Scale bar, 2 µm.

### MiniBAR promotes cilia growth and controls the local-ization of IFT88 and ARL13 into cilia

To investigate the function of MiniBAR, we used RNAi to knockdown Mini-BAR in RPE-1 cells (**Figure 4A**). Forty-eight hours after in-duction of ciliogenesis, the proportion of ciliated cells was modestly but reproducibly decreased upon MiniBAR deple-tion (**Figure 4B**). Importantly, cilia length was significantly reduced in cells harboring cilia (**Figure 4C**). Using stable cell lines that expressed siRNA-resistant version of wild type MiniBAR-GFP or mutants (**Figure S6B**), we observed that the percentage of ciliated cells as well as cilia length in knock-down cells were rescued by the expression of wild type MiniBAR (**Figure 4D-E**) but not MiniBARA140V or MiniBARA461R mutants (**Figure 4D-E**). Thus, MiniBAR promotes primary cilia formation and elongation in RPE-1 cells, and its interaction with both Rac1 and Rab35 is re-quired for its function in ciliogenesis.

**Fig. 4.**
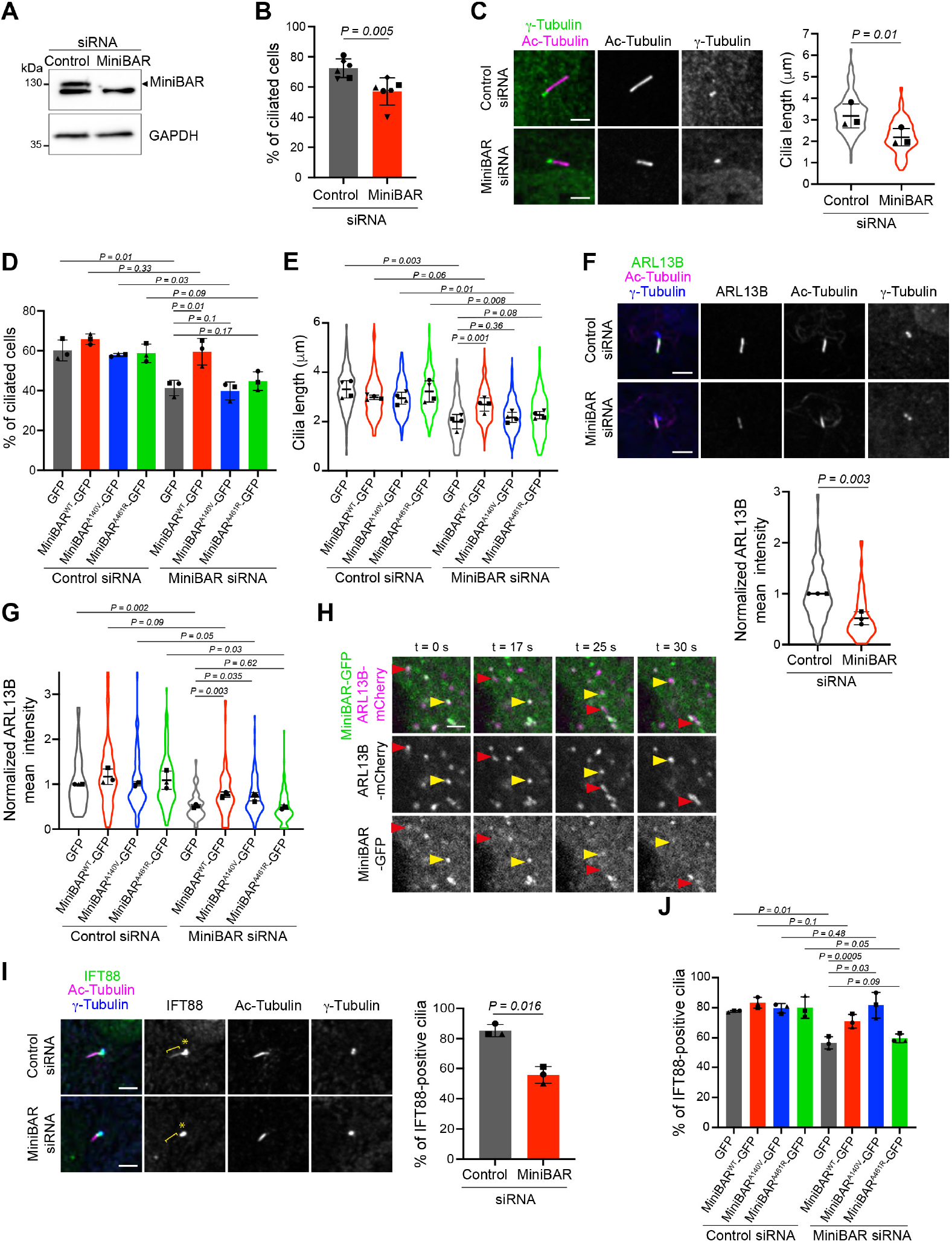
MiniBAR promotes cilia growth and controls the localization of IFT88 and ARL13 into cilia. (**A**) Lysates of RPE-1 cells transfected with control or MiniBAR siRNAs were blotted for MiniBAR and GAPDH (loading control).(**B**) Percentage of control-and MiniBAR-depleted cells with cilia (with acetylated-Tubulin-positive axonemes) measured 24 h after serum removal. Mean ± SD, N = 6 independent experiments. Paired student t-test.(**C**) Left panel: Representative images of cilia in control-and MiniBAR-depleted cells, 48 h after serum removal. Scale bars, 4 µm. Right panel: Cilia length based on acetylated-Tubulin staining in the aforementioned cells. Mean ± SD, n = 154-157 cells per condition, N = 3 independent experiments. Paired student t-test. In these and all subsequent "violin plots", the mean of each experiment as well as the distribution of all data points are indicated.(**D**) Percentage of cells with cilia measured 48 h after serum removal, in RPE-1 cells stably expressing either GFP alone, siRNA-resistant MiniBARWT-GFP, MiniBARA140V-GFP or MiniBARA461R-GFP and transfected with either control or MiniBAR siRNAs. Mean ± SD, n = 427-858 cells per condition, N = 3 independent experiments. Paired student t-tests. (**E**) Cilia length based on acetylated-Tubulin staining in the cells described in (D). Mean ± SD, n = 86-158 cells per condition, N = 3 independent experiments. Paired student t-tests.(**F**) Top panel: Representative images of cilia in control-and MiniBAR-depleted cells, 48 h after serum removal and stained for ARL13B. Scale bars, 4 µm. Bottom panel: mean ARL13B intensity in cilia in the aforementioned cells. Normalized mean ± SD, n = 144-154 cells per condition, N = 3 independent experiments. Paired student t-test.(**G**) Mean ARL13B intensity measured 48 h after serum removal, in RPE-1 cells described in (D). Mean ± SD, n = 90-130 cells per condition, N = 3 independent experiments. Paired student t-tests. (**H**) Snapshots of a movie of RPE-1 cells stably expressing MiniBAR-GFP, transfected with ARL13B-mCherry and incubated with SiR-Tubulin. Acquisition started 3-4 h after serum removal. Arrowheads: co-localization on two selected moving vesicles. Scale bar, 2 µm. (**I**) Left panel: Representative images of cilia in control-and MiniBAR-depleted cells, 48 h after serum removal, and stained for IFT88. Stars and brackets: ciliary base and the axoneme, respectively. Scale bars, 4 µm. Right panel: percentage of IFT88-positive cilia in the aforementioned cells. Mean ± SD, n = 114-128 cells per condition, N = 3 independent experiments. Paired student t-test.(**J**) Percentage of IFT88-positive cilia measured 48 h after serum removal, in RPE-1 cells described in (D). Mean ± SD, n = 85-210 cells per condition, N = 3 independent experiments. Paired student t-tests.

Both the Joubert syndrome protein ARL13B (associated with anterograde IFT-B trains) and the IFT-B train component IFT88 promote cilia elongation (Caspary et al., 2007; Cevik et al., 2013; Lu et al., 2015a; Nozaki et al., 2017; Yoder et al., 2002). Consistent with a key role of MiniBAR in controlling cilia length, we observed a 2-fold reduction of the mean intensity of endogenous ARL13B present in cilia upon MiniBAR depletion (**Figure 4F**). Expression of MiniBARWT or MiniBARA140V, but not of MiniBARA461R restored the cilia localization of ARL13B in MiniBAR-depleted cells (**Figure 4G**), indicating that MiniBAR’s interaction with Rab35 is key for ARL13B correct localization. ARL13B is known to traffic intracellularly on vesicles labelled by Rab22/ARF6 in non-ciliated cells (Barral et al., 2012). Since the ARF6 and the Rab35 pathways are physically and functionally connected (Allaire et al., 2013; Chesneau et al., 2012; Kobayashi et al., 2014a; Kobayashi et al., 2014b; Kobayashi and Fukuda, 2012; Rahajeng et al., 2012), we investigated whether ARL13B could partially traffic through the MiniBAR pathway. Time-lapse spinning disk confocal microscopy in cells co-expressing ARL13B-mCherry and MiniBAR-GFP revealed that ARL13B frequently co-localized with MiniBAR on moving vesicles (56 percent of ARL13B vesicles were positive for MiniBAR, n = 20-95 vesicles analyzed per cell from 6 cells, **Figure 4H and Movie 7**). We conclude that a major pool of ARL13B is transported by the MiniBAR vesicles in ciliated cells. In addition, the correct delivery of ARL13B in cilia relies on the presence of MiniBAR and on its interaction with Rab35.

Beside ARL13B, we observed that IFT88 was mislocalized in MiniBAR-depleted cells. In control cells, IFT88 localized both at the ciliary base (star) and within the cilium (bracket) in 85 percent of cilia (**Figure 4I**). In MiniBAR-depleted cells, IFT88 was detected only at the ciliary base (with an absence of IFT88 within cilium) more frequently than in control cells (**Figure 4I**). Again, the interaction between MiniBAR and Rab35 (but not with Rac1) was necessary for the correct localization of IFT88 within cilia, where it fulfills its IFT function (**Figure 4J**). We conclude that both MiniBAR and its interaction with Rab35 are required for the normal localization of ARL13B and IFT88 to cilia. This likely explains, at least in part, how MiniBAR controls cilia elongation.

### MiniBAR also controls cilia length by regulating actin-dependent contractility and RhoA vs. Rac1 activation

Beside trafficking and cargo delivery to the cilia, cilia elongation depends on the actin cytoskeleton, and excessive acto-myosin II activation is detrimental for ciliogenesis (Hoffman and Prekeris, 2022; Smith et al., 2020). Upon MiniBAR depletion, we noticed a clear difference in the global organization of the actin cytoskeleton. Instead of a few and thick stress fibers labelled with fluorescent phalloidin seen in control cells, numerous, closely packed and thin stress fibers were observed in MiniBAR-depleted cells (**Figure 5A**). Associated with these changes, the total F-actin levels and the Myosin II activation— were largely increased in MiniBAR-depleted cells (**Figure 5A-B**). Since MRLC phosphoryla-tion depends on kinases such as the Rho-activated kinase (ROCK), we hypothesized that the Rho GTPase might be over-activated in MiniBAR-depleted cells. Accordingly, pull-down assays using Rho effectors revealed that the levels of GTP-bound RhoA (RhoAGTP) were 3-fold higher upon MiniBAR depletion, as compared to control cells (**Figure 5C**). Consistent with the known mutual antagonism between Rac and Rho GTPases (Sander et al., 1999), Rac1GTP lev-els tended to be reduced in MiniBAR-depleted cells (**Figure 5C**). Thus, the presence of MiniBAR limits acto-myosin II-dependent cellular contractility.

**Fig. 5.**
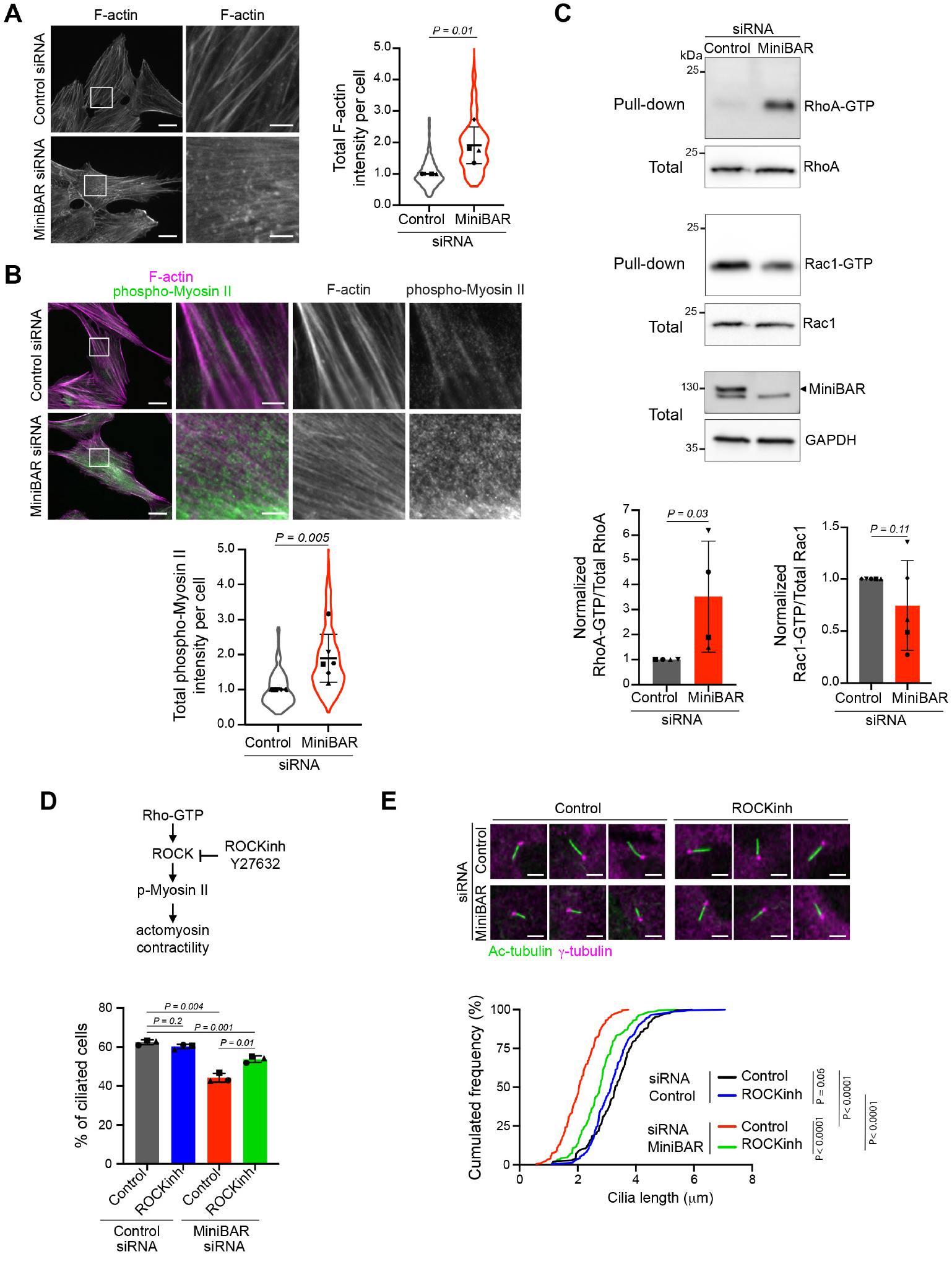
MiniBAR also controls cilia length by regulating actin-dependent contractility and RhoA vs. Rac1 activation. (**A**) Left panel: F-actin staining (phalloidin) in control-and MiniBAR-depleted cells, 48 h after serum removal. Scale bars, 20 µm (general view) and 4 µm (insets). Right panel: Total F-actin intensity in the aforementioned cells. Normalized mean ± SD, n = 263-402 cells per condition, N = 4 independent experiments. Unpaired student t-test. (**B**) Top panels: F-actin and phospho-S19 Myosin Regulatory Light Chain staining in cells described in (A). Scale bars, 20 µm (general views) and 4 µm (insets). Bottom panel: Total phospho-S19 Myosin Regulatory Light Chain intensity in the aforementioned cells. Normalized mean ± SD, n = 361-342 cells per condition, N = 5 independent experiments. Unpaired student t-test.(**C**) Top panels: Pull-down of endogenous RhoGTP and Rac1GTP from lysates of control-and MiniBAR-depleted RPE-1 cells, and revealed with anti-RhoA or anti-Rac1 antibodies, respectively. The total levels of endogenous RhoA, Rac1 and MiniBAR in each condition are also shown. Bottom panels: quantification of RhoAGTP over total RhoA and Rac1GTP over total Rac1. Normalized mean ± SD, N = 5 independent experiments. Unpaired student t-test.(**D**) Top panel: activation of myosin II contractility by the Rho/ROCK pathway. Bottom panel: Percentage of control-and MiniBAR-depleted cells with cilia measured 48 h after serum removal and treated or not with the ROCK inhibitor Y27632 for the last 24 h. Mean ± SD, n= 330-429 cells per condition, N = 3 independent experiments. Paired student t-tests. (**E**) Top panel: Representative images of cilia in cells described in (D). Scale bars, 4 µm. Bottom panel: Distribution of cilia length based on acetylated-Tubulin staining in the aforementioned cells. n = 187-219 cells per condition, N = 3 independent experiments. KS test.

To test whether the increased contractility observed upon MiniBAR depletion could impact on cilia length, we treated MiniBAR-depleted cells with the ROCK inhibitor Y27632 to reduce myosin II activity (**Figure 5D**). We found that both the decrease in the percentage of ciliated cells and the shortening of cilia observed upon MiniBAR depletion were partially rescued by treating the cells with Y27632 (**Figure 5D-E**). Of note, these relatively low doses of Y27632 had no impact on ciliogenesis in control cells suggesting a specific rescue of cilia defects in MiniBAR-depleted cells rather than a general promotion of ciliogenesis.

We conclude that MiniBAR limits the RhoAGTP /Rac1GTP balance, thus the cellular acto-myosin II-dependent contractility, which in turn favors normal ciliogenesis. Mechanistically, the ciliogenesis defects observed after MiniBAR depletion could be explained by a combined increase in cell contractility and defective trafficking or translocation of key cargos to cilia.

### MiniBAR depletion leads to dysfunctional cilia and hallmarks of ciliopathy in vivo

To investigate whether MiniBAR could play a role in ciliogenesis in vivo, we studied MiniBAR function during zebrafish embryogenesis. We identified a unique and highly conserved MiniBAR homologue in zebrafish (si:ch211-79l17.1 or Garre1) (**Figure S1A**). In situ hybridization showed that minibar mRNA was maternally expressed, then not detectable during gastrulation, and ubiquitously expressed at the 10-somite stage and 24 hpf (**Figure S7A**). Injection of antisense morpholinos (MO) targeting minibar mRNA at the 1-cell stage efficiently reduced MiniBAR expression at 24 hpf (**Figure 6A**) and resulted in multiple developmental defects characteristic of cilium dysfunction. Indeed, minibar morphants displayed a curved axis (**Figure 6B**), cystic kidneys (**Figure 6C**) and supra-numerous otoliths in the otic vesicles (**Figure 6D**). In addition, MiniBAR depletion resulted in left-right (L/R) asymmetry defects visible on heart jogging (**Figure 6E**) and pancreas positioning (**Figure S7B**). To further determine the role of MiniBAR in the establishment of L/R asymmetry, we analyzed the cascade of genes expressed asymmetrically in the left lateral plate mesoderm. In minibar morphants, the expression of the nodal related gene southpaw (spaw) and its downstream target pitx2 are altered, with higher frequency of embryos presenting bilateral, right or absent expression compared to control embryos (**Figure 6F and Figure S7C**). The striking heart L/R asymmetry defects observed upon MiniBAR depletion were partially rescued by co-injecting morpholino re-sistant miniBAR mRNA with the minibar MO (**Figure S7D**) and confirmed with a second, independent MO (MO minibar -2) and rescue experiments (**Figure S7E**). Long-term depletion of MiniBAR using maternal zygotic minibar -/- mutants led to the disappearance of full-length MiniBAR, as expected (**Figure S7F-G**), but we did not detect L/R patterning defects. This difference in phenotype between morphants and mutants could be due to a genetic compensation, a process described with several zebrafish mutants (El-Brolosy et al., 2019; Rossi et al., 2015). Nonetheless, CRISPR/Cas13d-mediated transient down-regulation (Kushawah et al., 2020) of MiniBAR after co-injection of specific minibar gRNAs and Cas13d at the 1-cell stage, resulted in MiniBAR knock-down and heart jogging defects 24 h later, similar to those observed upon MO injection (**Figure S7H**).

**Fig. 6.**
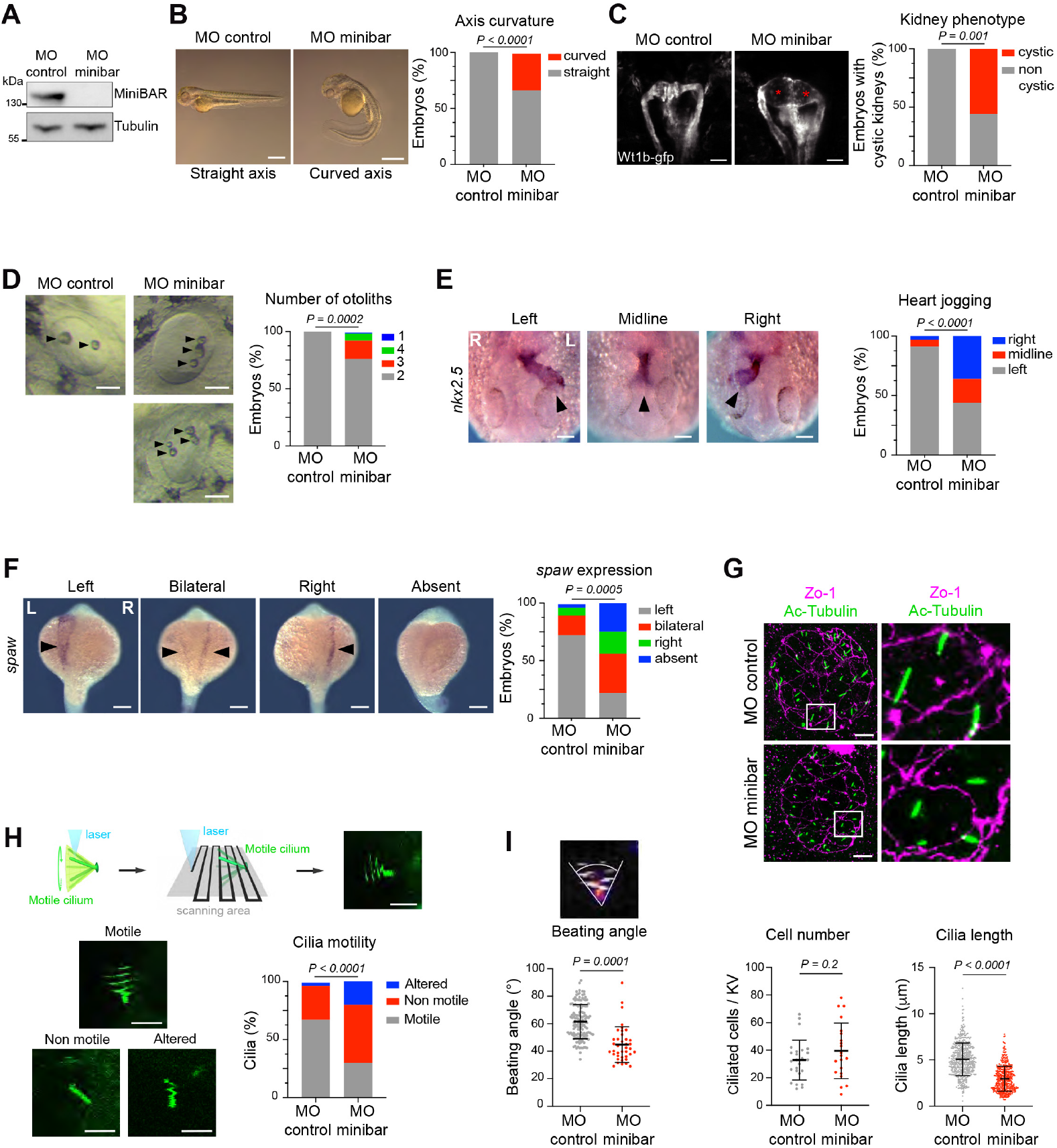
MiniBAR depletion leads to dysfunctional cilia and hallmarks of ciliopathy in vivo. (**A**) Lysates of 24 hours after fertilization (hpf) zebrafish embryos injected with either a control morpholino (MO control) or a morpholino targeting minibar (MO minibar) were blotted for MiniBAR and Tubulin (loading control). (**B**) Effect of minibar morpholino on axis curvature at 24 hpf. Scale bars, 500 µm. n = 40 and 92 embryos for control and minibar MO, respectively. Fisher’s Exact Test. (**C**) Effect of minibar morpholino on kidneys visualized with the wt1b:gfp line at 48 hpf. Asterisks indicate glomerular cysts. Scale bars, 50 µm. n = 14 and 18 embryos for control and minibar MO, respectively. Fisher’s Exact Test. (**D**) Effect of minibar morpholino on otoliths formation at 48 hpf. Scale bars, 50 µm. Arrowheads point to otoliths. n= 54 and 68 embryos for control and minibar MO, respectively. Fisher’s Exact Test. (**E)** minibar morphants exhibit heart jogging defects at 24 hpf, revealed by in situ hybridization for nkx2.5. Scale bars, 100 µm. n = 68 and 142 embryos for control and minibar MO, respectively. Chi-square test.(**F**) spaw expression at the 20-somite stage. Scale bars, 200 µm. n= 29 and 32 embryos for control and minibar MO, respectively. Fischer’s Exact Test. (**G**) Top panel: Zo-1 and acetylated-Tubulin immunostaining on Kupffer’s vesicle from 8 somite stage control and minibar morphants. Scale bars, 10 µm. Bottom left panel: Number of ciliated cells per Kupffer’s vesicle. n = 25 and 21 embryos for control and minibar MO, respectively. Unpaired student t-test. Bottom right panel: Quantification of cilia length. n = 19 embryos and 526 cilia for control and 14 embryos and 440 cilia for minibar MO. Linear mixed effect model. (**H**) Top panel: Imaging of a motile cilium with a point scanning technique leads to images with a characteristic pattern. Bottom left panel: Representative patterns obtained for normally beating, abnormally beating and immobile cilia, in βactin2:arl13b-gfp embryos. Scale bars, 5 µm. Bottom right panel: Quantification of the different beating patterns in control (159 cilia and 9 embryos) and minibar morphants (187 cilia and 9 embryos). Chi2 test. (**I**) Beating angle of motile cilia in control and minibar morphants. n = 9 embryos and 96 cilia for control and 9 embryos and 46 cilia for minibar MO, respectively. Linear mixed effect model.

L/R asymmetry patterning requires a break in symmetry that occurs in the zebrafish L/R organizer, the Kupffer’s vesicle (KV). Within the KV, cilia rotate and create a fluid flow that is critical to establish laterality (Grimes and Burdine, 2017). Cilia of the KV were shorter in minibar morphants compared to control embryos, while the number of ciliated cells per KV was unchanged (**Figure 6G**). The reduction in cilia length upon MiniBAR depletion was confirmed using CRISPR/Cas13d (**Figure S7H**). Using a two-photon micro-scope at a scanning speed lower than the cilia beat frequency, we quantified cilia motility in live transgenic embryos (Fer-reira et al., 2018) (**Figure 6H and Figure S7I**). We observed a higher proportion of immobile cilia in the KV of mini-bar morphants, compared to control embryos (**Figure 6H**). Furthermore, motile cilia displayed an abnormal beating pat-tern with a smaller angle of beating upon MiniBAR depletion (**Figure 6I**). Collectively, these results demonstrate a role of MiniBAR in cilia elongation in zebrafish, as seen in human RPE-1 cells. The importance of MiniBAR for normal cilia length and beating in the KV likely explains its requirement for the establishment of the L/R axis during zebrafish de-velopment. In addition, MiniBAR depletion leads to other characteristic defects observed upon inactivation of genes in-volved in ciliopathies.

## Discussion

Here, we report a conserved function in ciliogenesis of Mini-BAR, a unique dual Rac/Rab effector with a truncated BAR domain that controls both actin cytoskeleton contractility and membrane trafficking for cilia elongation.

### MiniBAR, the only known dual Rab/Rac effector with a truncated BAR domain

MiniBAR is one of the two reported dual Rac/Rab effectors. The only other example is Nischarin, which interacts with Rac1, Rab4A, Rab14 and Rab9A on overlapping domains, and regulates the maturation of early to late endosomes (Kuijl et al., 2013). Among all Rab GTPases tested, MiniBAR only interacts with active Rab35 (**Figure S1B**), suggesting that it is a specific Rab35 effector whereas most characterized Rab effectors interact with multiple Rab GTPases (Homma et al., 2021). Our biochemical data show that Rab35 and Rac1 can bind simultaneously to two separate domains of MiniBAR. The Rac1 binding domain overlaps with the domain of unknown function DUF4745 and forms a truncated BAR domain, which is not found in any other proteins. The Rab35BD central sub-domain adopts a Cystatin-like fold (**Figure S3C**) present in a number of proteins but the overall Rab35BD fold bears no similarity to other structurally resolved Rab effectors. Interestingly, careful examination of the presence of the Rac1BD and the Rab35BD across evolution revealed that the truncated BAR domain can be found in all Bilateria metazoa, i.e. in both Protostomia (such as Arthropods, Mollusks, Annelids, Worms) and Deuterostomia (such as Chordates including Vertebrates and Echinoderms) (**Figure S3A-C**). Furthermore, the DUF4745 (with putative Rac1BD) followed by the Rab35BD is a very ancient innovation and was likely present in the metazoa ancestor, which is known to have both Rho-and Rab-family proteins (**Figure S3D**).

### MiniBAR is present on intracellular vesicles and exhibit pulses along cilia

MiniBAR partially colocalizes with both Rac1 and Rab35 at the plasma membrane and on dynamic intracellular vesicles both in non-ciliated cells and during ciliogenesis (**Figure 2A-B, 3C-F**). MiniBAR crucially depends on its interaction with Rab35 to localize on vesicles (**Figure 2G**) and the co-overexpression of Rac1 and Rab35 induces the formation of intracellular MiniBAR tubules (**Figure 2D**), suggesting that both GTPases regulate MiniBAR activity on membranes. Altogether, MiniBAR is a novel component of the Rab35 endocytic recycling pathway. In addition to be transported on vesicles, MiniBAR and Rab35 exhibit striking, fast (15-30 seconds) pulses along cilia present 4- 8 hours after serum starvation (**Figure 3G,J**). Mechanistically, Rab35 pulses drive MiniBAR pulses (**Figure 3H**). To our knowledge, Rab35 and MiniBAR are the first membrane-associated proteins shown to pulse at the ciliary membrane. This raises two interesting questions that will require future investigations. First, how does Rab35/MiniBAR accumulate quickly and transiently at the membrane of cilia? The only case of transient accumulation of proteins in cilia that we are aware of has recently been described for the microtubule-associated kinesin KIF13b and is closely coordinated with IFTs, although the exact mechanism of transient accumulation is not yet understood (Juhl et al., 2023). Rab35- MiniBAR might be associated with IFTs but this hypothesis is difficult to reconcile with the observation that MiniBAR signals spread in a wave-like manner from either tip to base (**Movie 5, time 98 s**), center to both base and tip (**Movie 5, time 180 s**) or base to tip (**Movie 6, time 100 s**). An interesting possibility is that a soluble pool (Breslow et al., 2013) of GDP-Rab35 is locally activated within cilia and thereby concentrates at the ciliary membrane, where it recruits MiniBAR. We cannot exclude that Rab35/MiniBAR complexes may pulse at the surface of intracellular ciliary vesicles, but we consider this unlikely since at 4-8 hours post-starvation cilia were all (n = 25/ 25) labelled within 2-5 min upon addition in the extracellular medium of CellBrite® (a fluorescent lipid that integrates the plasma membrane and that shows no sign of internalization at this time scale). Interestingly, in-traciliary pulses of Ca2+ have been reported in the mouse node and the zebrafish KV (Mizuno et al., 2020; Yuan et al., 2015), with similar frequency in mouse embryos (1 spike per min) and contribute to L/R asymmetry establishment in vivo (Mizuno et al., 2020). Ca2+ pulses might control Rab35 GEF/GAP activities and consequently MiniBAR pulses. A second question raised by our observations is the physiological relevance of MiniBAR pulses for cilia growth. Since MiniBAR pulses during the first 24 hours upon cilia induction but not later, when cilia do not grow any longer in our experimental conditions, one can speculate that it might help to deliver specific cargos during the early steps of ciliogenesis when cilia elongate.

### MiniBAR promotes cilia elongation by controlling membrane trafficking of key cilia proteins and by limiting ac-to-myosin-dependent contractility

Both in human RPE-1 cells and in the KV cells in zebrafish embryos, MiniBAR depletion leads to abnormally short cilia (**Figure 4C, 6G**) indicating that MiniBAR is a rate limiting factor for cilia growth. Point mutations that selectively disrupt either the binding with active Rac1 or with active Rab35 demonstrate that the interaction with each GTPase is required for normal ciliogenesis (**Figure 4D-E**). Mechanistically, our data indicate that MiniBAR promotes ciliogenesis by controlling two pathways (**Figure 7**).

**Fig. 7.**
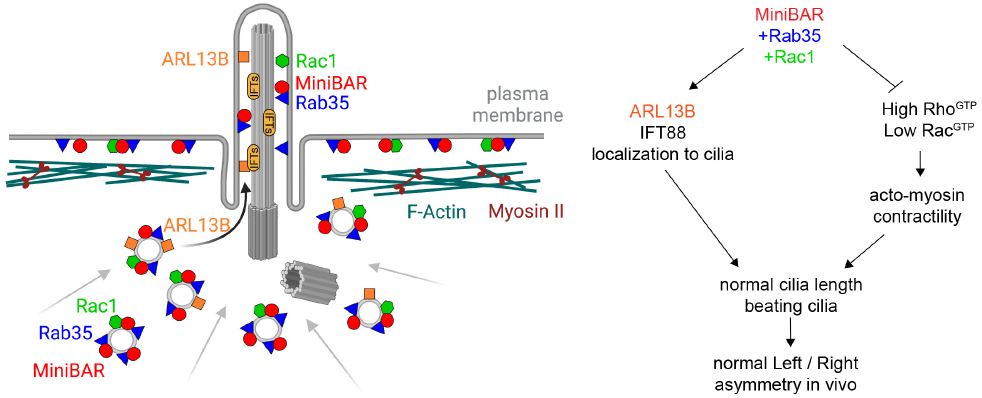
Summary model of the functional role of MiniBAR in ciliogenesis by controlling membrane trafficking and acto-myosin contractility. Left panel: Model representing the localization of MiniBAR (red), Rab35 (blue) and Rac1 (green) at the plasma membrane and on intracellular vesicles at the ciliary base, and the role of MiniBAR in the localization of ARL13B (orange) and IFT88 (not represented) in cilia. Right panel: summary model of the functional role of MiniBAR in interaction with Rab35 and Rac1 for promoting cargo delivery in cilia and limiting acto-myosin contractility, both necessary for correct cilia length and beating, and thus normal left-right asymmetry in vivo.

First, MiniBAR helps to deliver at least two proteins important for ciliogenesis within cilia: ARL13B and IFT88 (**Figure 7, right panel**). ARL13B is well known to promote cilia elongation (Caspary et al., 2007; Cevik et al., 2013; Lu et al., 2015a; Nozaki et al., 2017) but how it is targeted to the cilia is not well understood. Here, we found that a pool of ARL13B is transported on dynamic MiniBAR vesicles (**Figure 4H**), which are actively trafficked toward the ciliary base throughout ciliogenesis (**Figure 3B-F**). This suggests that MiniBAR might play a role both for the ciliary vesicle formation/maturation and cilia elongation by delivering key cargos to growing cilia, such as ARL13B. Accordingly, the accumulation of ARL13B within cilia depends on MiniBAR and on its interaction with RaB35 (**Figure 3G,H**). Of note, Rab35 likely plays multiple roles in ARL13B trafficking in and out of cilia, possibly via multiple effectors, since Rab35 restricts ciliary accumulation of ARL13B (Kuhns et al., 2019). A strength of our study is to unveil the specific role of Rab35 with MiniBAR, thanks to the crystal structure-guided design of a point mutant that selectively abolishes the interaction between Rab35 and MiniBAR while leaving untouched the interaction of Rab35 with its other effectors. The localization within cilia of the anterograde IFT component IFT88 is also perturbed in MiniBAR-depleted cells and our data suggest that MiniBAR and its interaction with Rab35 promotes either the translocation of IFT88 into cilia or its delivery to the ciliary base (**Figure 4I-J**). Altogether, the Rab35/MiniBAR trafficking pathway helps to deliver two limiting factors for cilia growth, which can explain MiniBAR’s requirement for cilia elongation. Besides regulating trafficking, MiniBAR limits actomyosin-dependent contractility (**Figure 5A-C and Figure 7, right panel**), which in turn favors cilia elongation. Consistently, the cilia length defects found in MiniBAR-depleted cells could be largely rescued by inhibiting ROCK, a downstream Rho effector that activates myosin II (**Figure 5E**). The regulation of actomyosin-dependent contractility likely involves the pool of MiniBAR at the plasma membrane (**Figure 2A-B**) rather than on internal vesicles. However, how MiniBAR controls the RhoGTP/RacGTP balance is unknown, but MiniBAR could act directly, or together with yet to be discovered partners. Interestingly, the fact that reducing myosin II activity does not fully rescue cilia length suggests that MiniBAR controls both pathways —trafficking and contractility. It should be noted that increased levels of stress fibers and hyperactivation of RhoA have been also observed upon mutation/inactivation of different genes involved in ciliopathies (Hoffman and Prekeris, 2022; Smith et al., 2020). Furthermore, ROCK inhibition can rescue ciliary defects in several ciliopathy models including Bardet-Biedl syndrome and polycystic-kidney disease models (Cai et al., 2018; Hernandez-Hernandez et al., 2013; Kim et al., 2015; Stewart et al., 2016; Streets et al., 2020), as observed upon MiniBAR depletion. While it is well established that membrane traffic and cell contractility contribute to ciliogenesis, little is known about proteins that regulate both. It was recently reported that i) Rab19 helps to clear the actin cortex at the ciliary base —through unknown mechanisms— and promote cilia formation (Jewett et al., 2021) and ii) the depletion of the dual Rab10/RhoA GEF DENN2B increases the number of ciliated cells and cilia length (Kumar et al., 2022)—the opposite of what was observed upon MiniBAR depletion. Thus, there is increasing evidence that cell contractility and membrane traffic are intimately linked and must be tightly regulated during ciliogenesis. We propose that MiniBAR helps to coordinate these two processes to control cilia elongation.

### MiniBAR controls cilia length and L/R asymmetry in vivo

Both Rac1 and Rab35 GTPases were independently shown to promote ciliogenesis and their disruption leads to L/R asymmetry defects in zebrafish, but direct effectors are yet to be discovered (Epting et al., 2015; Hashimoto et al., 2010; Kim et al., 2010; Kuhns et al., 2019). MiniBAR might thus represent a missing link between these two GTPases. MiniBAR-depleted zebrafish embryos display several typical defects associated with ciliopathies, such as cystic kidneys and situs inversus (**Figure 6A-F and S7B**). As in RPE-1 cells, cilia are shorter in the KV and the proportion of non-beating and abnormally beating cilia were increased (**Figure 6G-I**). This likely explains the defective bias in the expression of asymmetrically expressed transcription factors (**Figure 6F and S7C**). Other defects are likely due to shorter/defective cilia in other locations, consistent with the ubiquitous expression of minibar mRNA in the embryo (**Figure S7A**). Altogether, the in vivo data raises the possibility that genetic ciliopathies of unknown origin in Humans might originate from mutations in MiniBAR.

MiniBAR is likely important also in non-ciliated cells since active MiniBAR-positive vesicles are found in proliferating cells (**Figure 2**). Interestingly, the three mutations shown to disrupt the interaction between MiniBAR and Rac1, A140V, 140T and M144K (**Figure S5A**) are somatic mutations found in cancer biopsies of endometrioid carcinoma, B cell lymphoma, lung squamous cell carcinoma and intestinal adeno-carcinoma —in which inactivating RhoA mutations are often observed (Tate et al., 2019; Wang et al., 2014). Thus, MiniBAR may represent a key integrator of Rac/Rab signaling that controls trafficking and the actomyosin cytoskeleton in ciliogenesis and beyond, notably in pathological situations such as tumorigenesis.

## Methods

### Plasmids

pDEST 3xFLAG C1-Rab35WT, pDEST 3xFLAG C1-Rab35Q67L and pDEST 3xFLAG C1-Rab35S22N were described in (Klinkert et al., 2016). pmCherryFP-Rab35 WT was described in (Cauvin et al., 2016). pcDNA FLAG-Rac1WT, pcDNA FLAG-Rac1Q61L, pcDNA FLAG-Rac1T17N, peGFP-Rac1 WT and pcDNAm FRT PC mCherry Rac1 WT were kind gifts from Alexis Gautreau. pARL13B-mCherryFP was a kind gift from Alexandre Benmerah. Yeast two hybrid vectors pLexA and pGAD have been described in as described in (Klinkert et al., 2016). Human MiniBAR cDNA (KIAA0355) was subcloned into Gateway pENTR plasmid and -LAP-GFP, -LAP-mCherry and -iRFP expression vectors were generated by LR recombinaison (Thermo Fisher). Point mutations in MiniBAR have been generated using NEBaseChanger (NEB), were amplified by PCR and introduced into pENTR gateway vectors, then recombined into the transient expression vector (-LAP-GFP, -LAP-mCherry, GAD) or lentiviral-GFP destination vector (ThermoFisher scientific). For rescue experiments, siRNA-resistant versions have been obtained by mutating 6 bp of the siRNA-targeting sequence using Quickchange (Agilent).

### Antibodies

The following antibodies were used for immunofluorescence in human cells: Mouse anti-G-Tubulin (Sigma T6557, 1:1000, Methanol fixation in Figure 3B, 4C,I and 1:200, PFA fixation in Figure 3A, 4F, 5E); Human anti-Acetylated-Tubulin (Institut Curie: Therapeutic recombinant antibodies platform clone (C3B9-hFc), 1:200, PFA fixation in Figure 3A, 4F, 5E and methanol fixation in Figure 4C,I), Rabbit anti-pMRLC (Cell signaling 3671, 1:100, PFA fixation), Rabbit anti-ARL13B (Proteintech 17711-1-AP, 1:200), Rabbit anti-IFT88 (kind gift from Chantal Desdouets (Robert et al., 2007), 1:50, methanol fixation), Rabbit anti-DCP1 (Abnova 55802, 1:200, PFA fixation), Rabbit anti-GW182 (Abcam ab70522, 1:200, PFA fixation). Rabbits were immunized for 72 days by Covalab (Bron, France) with purified human MiniBAR556-971 to obtain antisera directed against the C-terminal half of MiniBAR. Rabbit anti-MiniBAR (1:1000, methanol fixation). Fluorescently Alexa-488, Cya3, Cya5 labelled secondary antibodies were purchased from Jackson Laboratories (1:500). Phalloidin 647 was purchased from Cell signaling Technology and CellBrite® was purchased from Biotium (30107, 1:2000). The following antibodies were used for whole mount staining in zebrafish embryos: Zo-1 (Invitrogen 33-9100, 1:250), Acetylated tubulin (T6793, 1:500), anti-mouse IgG2b Alexa Fluor 488 (Invitrogen A21141, 1:500) and anti-mouse IgG1 Alexa Fluor 594 (Invitrogen A21125, 1:500). The following antibodies were used for western blots of human cell extracts: Mouse anti-FLAG (Sigma F1804, 1:1000); Mouse anti-GAPDH (Proteintech 60004-1-Ig, 1:40 000); Mouse anti-RhoA (Cytoskeleton ARH05, 1:500); Mouse anti-Rac1 (Cytoskeleton ARC03, 1:500); Mouse anti-G-Tubulin (Sigma T4026, 1:5000); Rabbit anti-MiniBAR (1:500). Secondary horseradish-peroxidase-coupled antibodies were purchased from Jackson Laboratories.

### Protein expression, purification

The human constructs (Rac1, Rab35, Rac1BD (MiniBAR70-208), Rab35BD (MiniBAR225-553) and Rac1BD-Rab35BD (MiniBAR70-553), were cloned in pPROEX-HTb or pRSFduet plasmids with N-terminal His-tag and rTEV cleavage sites; expressed in E. coli BL21(DE3) or BL21(DE3)RIPL cells. The cells were grown in 2xYT media to 0.6 OD600 a 37°C, induced with 0.2 mM IPTG at grown overnight at 17 °C. The cells were harvested by centrifugation, disrupted by sonication in (50 mM Tris pH8, 150 mM NaCl, 2 mM MgCl2, 1 mM TCEP, 1 mM PMSF, complete protease inhibitor mix (GE Healthcare), 5 percent glycerol (v/v), 40 mM imidazole). The cell lysate was applied on Ni-NTA column (GE Healthcare) after clarification by centrifugation. The Ni-NTA column was washed with (50 mM Tris pH8, 150 mM NaCl, 2 mM MgCl2, 1 mM TCEP, 5 percentglycerol, 40 mM imidazole) and the protein was eluted with 250mM imidazole. The His-tag was removed by overnight incubation with rTEV protease (1:100 protein:protease ratio). Gel-filtration was performed as a final purification step, using Superdex 200 or Superdex 75 columns (GE Healthcare) in running buffer: 50 mM Tris pH8, 50 mM NaCl2, 2 mM MgCl2, 2 mM TCEP.

The proteins were concentrated, flash frozen in liquid nitrogen and stored at -80 °C. The active GTP-bound form of Rac1 was produced by introducing Q61L mutation (Rac1Q61L). The active form of Rab35 was generated by GPPNHP nucleotide exchange before the final gel-filtration: the purified Rab35 protein was incubated with 10 times molar excess of GPPNHP in 10 mM EDTA containing buffer, the exchange was terminated by adding 20 mM MgCl2.

### ITC

Isothermal titration calorimetry experiments were carried out on a MicroCal ITC-200 titration microcalorimeter (Malvern) at 10 °C. The protein samples were in 50 mM Tris pH 8, 50 mM NaCl, 2 mM MgCl2, 2 mM TCEP. The titration processes were performed by injecting 2 µL aliquots of protein samples from a syringe (active form of Rac1 or Rab35 at a concentration of 480 µM) into protein samples in cell (MiniBAR fragments at a concentration of 60 µM) at time intervals of 2 min to ensure that the titration peak returned to the baseline. The titration data were analyzed using the instrument producer provided software (Origin7 (Edwards, 2002)) and fitted by the one-site binding model.

### Analytical SEC and SEC-MALS

Absolute molar masses of proteins were determined using size-exclusion chromatography combined with multi-angle light scattering (SEC–MALS). Protein samples (Rac1BD, Rab35BD and Rac1BD-Rab35BD) (40 µL; 10-25 mg/ml) were loaded onto a Superdex-200 increase 5/150 column (GE Healthcare) or BioSec5 300 (Agilent) or XBridge BEH SEC 200 A (Waters) in 50 mM Tris pH 8, 50 mM NaCl, 2 mM MgCl2, 2 mM TCEP, at 0.3-0.5 ml/min flow rate using a Dionex UltiMate 3000 HPLC system. The column output was fed into a DAWN HELEOS II MALS detector (Wyatt Technology). Data were collected and analyzed using ASTRA software (Wyatt Technology). Molecular masses were calculated across eluted protein peaks. The eluted protein peak fractions were collected and analyzed by SDS-PAGE. Analytical SEC was performed using the same setup excluding MALS detector.

### Crystallization and X-ray structure determination

Crystals were obtained by the hanging drop vapor diffusion method at 17 °C by mixing of 1 µL of concentrated protein solution with 1 µL of crystallization solution. Initial crystals were improved by microseeding (Bergfors, 2003). Crystals of the selenomethionine substituted Rab35BD were grown with 1.26 M Sodium Malonate pH 6.3, 5 percent (v/v) glycerol, 5 percent ethylene glycol, 10 mM DTT. The crystals were cryoprotected using the crystallization solution with increased Sodium Malonate concentration to 3M before flash freezing in liquid nitrogen. A single-wavelength anomalous diffraction dataset was collected at the SOLEIL synchrotron Proxima-1 beamline, processed with XDSME package (Kabsch, 2010; Legrand, 2017) and phased using SHELX program suite (Sheldrick, 2008). The phases were improved using Phaser (Read and McCoy, 2011) with subsequent density modification using Pirate (Cowtan, 2000). The initial protein model was built with Buccaneer (Cowtan, 2006) and refined with BUSTER (Bricogne G. and Roversi P, 2017). After iterative rebuilding in COOT (Emsley and Cowtan, 2004) and refinement with PHENIX (Adams et al., 2010) using the data set reprocessed with AutoPROC (Evans, 2006; Evans and Murshudov, 2013; Kabsch, 2010; Vonrhein et al., 2011; Winn et al., 2011) the final model (Suppl. Table 1) was deposited to the Protein Data Bank (PDB). Rac1BD was crystallized in 100mM Ammonium Citrate dibasic, 16 percent (w/v) PEG 3350, The crystals were cryoprotected by supplementing the crystallization solution with 22 percent (v/v) ethylene glycol. A native data set was collected at the SOLEIL synchrotron Proxima-2 beamline. The data set was processed with AutoPROC (Vonrhein et al., 2011) and the structure was determined by molecular replacement using Archimboldo program (Rodríguez et al., 2012) from CCP4 program sute (Winn et al., 2011), model rebuilding was performed using COOT (Emsley and Cowtan, 2004) and refinement -using PHENIX (Adams et al., 2010), the final structure (Suppl. Table 1) was deposited to the PDB. All structure figures were prepared using PyMOL (Schrödinger and DeLano, 2020).

### Small angle X-ray scattering (SAXS)

SAXS data collection were performed at SOLEIL synchrotron SWING beam-line (Suppl. Table 2), the samples were subjected to on-line size exclusion purification with the HPLC system connected to a quartz capillary placed under a vacuum cell (Adams et al., 2010). Data reduction, averaging of identical frames corresponding to the elution peak and buffer subtraction were performed with the SWING in-house software Foxtrot. The radius of gyration (Rg) and forward intensity at zero angle I(0) were derived by the Guinier approximation using the software PRIMUS (Konarev PV et al., 2003). The maximum dimension (Dmax) and the Rg were derived from the pair distribution function P(r), calculated with the software GNOM (Svergun DI, 1992). The molecular weight was calculated with the PRIMUS Molecular Weight wizard (Konarev PV et al., 2003). The MultiFoXS web-server (Schneidman-Duhovny et al., 2016) was used to fit the theoretical scattering intensity from the X-ray structure into the experimental SAXS data (Suppl. Table 2); side chains and loops missing in the X-ray structures were modelled using Modeller (Webb and Sali, 2016) before the fitting.

### Cell culture

hTERT RPE-1 cells (hTERT-immortalized retinal pigment epithelial cells, ATCC CRL-4000™ a kind gift of A. Benmerah) were grown in DMEM-F12 medium (Gibco) supplemented with 10 percent fetal bovine serum (FBS; Dutscher, 500105R1) and 1 percent penicillin/streptomycin (Gibco, 15140122) in 5 percent CO2 at 37 °C. For serum starvation experiments, cells were washed twice with DMEM-F12 without SVF and incubated with DMEM-F12 without SVF. For experiments with the ROCK inhibitor (Figure 5E), RPE-1 cells transfected with either siRNAs targeting Luciferase or MiniBAR for 72 h were treated with Y27632 (Calbiochem 146986-50-7) at 50 µM for 24 h in DMEM-F12 without SVF before fixation.

### Stable cell lines

Stable cell lines expressing low levels of siRNA-resistant, -GFP-tagged, full-length MiniBARWT and mutants (MiniBARA140V or MiniBARA461R) were generated by lentiviral transduction in RPE-1 cells. Briefly, lentiviral particles were produced in the HEK293 FT packaging cells transfected with MiniBAR constructs and the Excelenti LTX Lentivirus Packaging mix (Oxford Genetics) using lipofectamine following manufacturer’s instructions. After 48 h, the HEK293 FT culture supernatants were added to RPE-1 cells for 24 h. A week later, GFP-positive cells were sorted by FACS.

### Zebrafish strains

Embryos were obtained by natural spawning of AB, Tg(wt1b:EGFP) (Bollig et al., 2009) (Bollig et al. 2009), Tg(actb2:Mmu.Arl13b-GFP) (Borovina et al., 2010) and minibarsa35606 adult fishes. Embryos were incubated at 28°C and staged in hours post-fertilization (hpf) as described (Kimmel et al., 1995). All animal studies were approved by the Ethical Committee N°59 and the Min-istère de l’Education Nationale, de l’Enseignement Supérieur et de la Recherche under the file number APAFIS15859- 2018051710341011v3. minibar sa35606 mutant minibarsa35606 mutants carry a point mutation at Chr 18: 369582 location, leading to a premature stop codon. The line was obtained from the Zebrafish International Resource Center (ZIRC). Fishes were genotyped by fin clipping. DNA preparation was performed by placing fin clip in lysis buffer (50 mM KCl, 10 mM Tris pH 8.3, 0.3 percent Tween-20, 0.3 percent NP-40) with fresh Proteinase K (Merck Cat 03115887001) at 55 °C overnight followed by 15 min of inactivation of proteinase K inactivated at 95 °C. Genotyping was performed by PCR followed by restriction enzyme digestion. The following PCR primers were used: Sa35606 Fwd-5’- GACCATAGGCTCCTCCCCTTCTGG-3’ Sa35606 Rev-5’- CCATGACATGTTCCTGCACGTCCTTCAGCT-3’. The PCR product (218 bp spanning the mutation) was digested with PvuII. Digestion can only occur in WT amplification, leading to a 190 bp fragment in WT and 218 bp in mutants.

### Yeast two hybrid assays and screens

A placenta random-primed cDNA library was screened by Hybrigenics Services SAS using human LexA-RAB35Q67L (aa 1-194, to delete the terminal prenylated cysteines) as a bait in the presence of 2 mM 3-aminoTriazole. 26 out of 227 isolated clones from 108 million screened interactions encoded part of KIAA0355. The same library was screened with human LexA-Rac1G12V, C189S (active Rac1 with mutated terminal prenylated cysteine). 336 clones were selected from 60 million interactions tested, of which 10 encoded for KIAA0355 fragments. Furthermore, 8 clones encoding for KIAA0355 fragments were also isolated using a Rac1G12V, C189S - LexA fusion as a bait, from 53 million interactions tested and a total of 153 selected clones. No 3-aminotriazole was used in the Rac1 screens. Yeast two-hybrid experiments were performed by co-transforming the Saccharomyces cerevisiae reporter strain L40 with either pGAD-MiniBAR full length, MiniBAR Rac1BD, or MiniBAR Rab35BD together with either pLex-human Rab35WT, pLex-human Rab35Q67L, pLex-human Rab35S22N, or pLex-human Rac1WT, pLex-human Rac1Q61L, pLex-human Rac1T17N. Transformed yeast colonies were selected on DOB agarose plates without Tryptophane and Leucine. Colonies were picked and grown on DOB agar plates with Histidine to select co-transformants and without Histidine to detect interactions, as described in (Fremont et al., 2017).

### Human cell transfection and RNA interference

Plasmids were transfected in RPE-1 cells for 24 h (Figure 2B,C; Figure 3D,F,J) or 48 h (for Figure 4H) using X-tremeGENE 9 DNA reagent (Roche) following the manufacturer’s instructions. For Rab35 silencing experiment, MiniBAR-GFP stable cell line was transfected with 25 nM of siRNAs targeting luciferase (5’-CGUACGCGGAAUACUUCGA-3’) or human Rab35 (5’-GCUCACGAAGAACAGUAAA-3’) using Lipofectamine RNAiMAX (Invitrogen), following the manufacturer’s instructions. Time lapse microscopy after siRNA Rab35 (Figure 3H) were started 72 h after siRNA transfection. For MiniBAR silencing experiments, RPE-1 cells or MiniBAR-GFP stable cell lines were transfected with 25 nM of siRNAs targeting either luciferase or human MiniBAR (5’-CUGCAAAUUUUACGGAUCA-3’) using Lipofectamine RNAiMAX (Invitrogen), following the manufacturer’s instructions. For MiniBAR siRNA experiments with serum removal (Figure 4A-G, 4I-J and 5A-E), the serum was removed to induce ciliogenesis for 48 h after 24 h of siRNA transfection.

### Western blots

RPE-1 cells transfected with siRNAs were lysed in cell lysis buffer (cytoskeleton, Part CLB01) supplemented with protease inhibitors. 30 µg of lysate was migrated in 4-15 percent gradient SDS–PAGE gels (BioRad Laboratories), transferred onto PVDF membranes (Millipore) and incubated with indicated primary antibodies in TBS, 5 percent low-fat milk and 0.1 percent Tween20. The membranes were incubated with HRP-coupled secondary antibodies and revealed by chemiluminescence (GE Healthcare). MiniBAR-GFP stable cell lines transfected with siRNAs were directly collected in 1x loading buffer, boiled 5 min and 10 µL of total extracts were migrated in SDS-PAGE. 3 to 5 zebrafish embryos were directly collected in 1x loading buffer, boiled 5 min and 1/4 of the total extracts was migrated in SDS-PAGE.

### Pull down assay

RhoAGTP and Rac1GTP pull down assays were performed using the RhoA and Rac1 Activation Assay Biochem Kit (Cytoskeleton BK030) according to the manufacturer’s instructions.

### Co-immunoprecipitation assays

RPE-1 cells were transfected with the different Flag constructs for 48 h using X-tremeGENE 9 DNA reagent (Roche). Cells were lysed in 25mM Tris (pH 7.5), 150mM NaCl, 10mM MgCl2 and 0.1 percent Triton X-100. Supernatants (10min at 20 000g) were incubated with anti-Flag M2 affinity beads (Sigma-Aldrich A2220) for 3h, washed with lysis buffer and re-suspended into 1X Laemmli buffer and boiled at 95°C for 5min. The amount of co-immunoprecipitated MiniBAR endogenous protein in each condition was probed by western blot using anti-MiniBAR antibodies.

### Immunofluorescence in human cells and image acquisition

Cells were grown on coverslips and then fixed with 4 percent paraformaldehyde (PFA) for 15 min at room temperature or with ice-cold methanol for 3 min at -20 °C depending on the antibodies (see Antibodies section). For ciliogenesis induction, the serum was removed 48 h before fixation. For ARL13B and IFT88, cells were permeabilized after fixation with 0.1 percent Triton-X100 for 3 min, blocked with 0.2 percent BSA/PBS for 30 min and successively incubated for 1 h at room temperature with primary and secondary antibodies (1:500; Jackson ImmunoResearch) diluted in PBS containing 0.2 percent BSA. For phospho-myosin II and phalloidin (Thermo fisher scientific, 1/200 PFA fixation) staining, cells were fixed with 4 percent paraformaldehyde (PFA) in cytoskeleton buffer (10 mM MES, 138 mM KCl, 3 mM MgCl2, 2 mM EGTA supplemented with sucrose) permeabilized with 0.1 percent Triton-X100 for 5 min and incubated with primary antibodies and phalloidin (1:200) overnight in cytoskeleton buffer containing 0.2 percent BSA. For MiniBAR staining, cells were permeabilized and saturated with 0.2 percent BSA/PBS/saponin 0.1 percent for 30 min, then incubated with primary and secondary antibodies (1:500; Jackson ImmunoResearch) diluted in 0.2 percent BSA/PBS/saponin 0.1percent. Cells were mounted in Mowiol (Calbiochem) after DAPI staining (0.5 µg /mL, Serva). Images in Figure 2G, 3B and 4A-B were acquired with an inverted TiE Nikon microscope, using a x100 1.4 NA PL-APO objective lens or a x60 1.4 NA PL-APO VC objective lens and MetaMorph software (MDS) driving a CCD camera (Photometrics Coolsnap HQ). For co-transfected cells (Figure2G), cells were fixed 24 h after transfection and directly mounted in Mowiol (Calbiochem). Images in Figure 4C, 4F, 4I and 5E were acquired using an inverted Nikon Eclipse Ti-E microscope equipped with a CSU-X1 spinning disk confocal scanning unit (Yokogawa) and with an electron-multiplying CCD (EMCCD) camera (Evolve 512 Delta, Photometrics) or a Prime 95S sCMOS camera (Teledyne Photometrics).

SIM was performed on a Zeiss LSM 780 Elyra PS1 microscope (Carl Zeiss, Germany) using C Plan-Apochromat 63×/1.4 oil objective with a 1.518 refractive index oil (Carl Zeiss). The fluorescence signal was detected on an EMCCD Andor Ixon 887 1K. Raw images were composed of fifteen images per plane per channel (five phases, three angles), and acquired with a Z-distance of 110 nm. The SIMcheck plugin (Ball et al., 2015) in ImageJ was used to analyze the quality of the acquisition and the processing in order to optimize parameters for resolution, signal-to-noise ratio, and reconstruc-tion pattern.

For spinning disk confocal experiments, RPE-1 cells or MiniBAR-GFP stable cell lines were plated on 35 mm glass bottom plates (MatTek). For transfection experiments, cells were imaged 24 h (Figure 2B-C and 3D,F,J) or 48 h after transfection (Figure 4H). For serum starvation experiment, serum was removed 3-4 h before imaging and SiR-Tubulin was added at the same time and kept during the time of the movie for 2-4 h. Cells were incubated in an open chamber (Life Imaging) equilibrated with 5 percent CO2 and main-tained at 37 °C. Images were acquired using an inverted Nikon Eclipse Ti-E microscope equipped with a CSU-X1 spinning disk confocal scanning unit (Yokogawa) and with an electron-multiplying CCD (EMCCD) camera (Evolve 512 Delta,Photometrics) or a Prime 95S sCMOS camera (Teledyne Photometrics).

### Image quantification and kymograph analysis

Quantification of MiniBAR mean intensity around centrosomal area was done manually following MiniBAR staining around G-Tubulin staining using Fiji. For quantification of total F-actin and mean phospho-myosin per cell, cell contours were defined manually. Because of the cell and the cilium movements, a kymograph must be extracted from end-points that follow the cilium base and tip as they move. To this end we used TrackMate (Ershov et al., 2022) to manually track the cilium base and tip. For each time-point, we then extracted an intensity profile between these two points, averaged over a width of 5 pixels. The intensity profiles are collected for every frame of the movie, following the cilium movement. The profiles are then stacked along time and centered in their middle to generate a kymograph like the one shown in Figure 3g. Imaging of zebrafish embryos Imaging of Kupffer’s vesicle and kidney was done on an inverted TCS SP8 confocal microscope (Leica) using a High NA oil immersion objective (HC PL APO 63x 1.40, Leica). Whole-mount immunostained KV were dissected and mounted on a slide. Imaging of Kupffer’s vesicle for cilia motility quantification was done under an upright TriM Scope II (La Vision Biotech) two-photon microscope equipped with an environmental chamber (okolab) at 28 °C. Injected embryos were mounted at 8- somite in 0.2 percent agarose in embryo medium. Whole embryo imaging was performed with a M205FCA stereomicroscope (Leica) and a MC170HD Camera (Leica). Images were processed in Fiji and Adobe Photoshop.

### Whole-mount In Situ Hybridization in zebrafish embryos

Zebrafish embryos were fixed in 4 percent paraformaldehyde at 4 °C for 48 hpf. Whole-mount in situ hybridisations were performed as previously described (Hauptmann and Gerster, 1994) using spaw, ptx2, nkx2.5, insulin, and minibar probe.

### Whole-mount immunostaining in zebrafish embryos

Embryos from 8 to 10-somite stage were fixed with 4 percent paraformaldehyde overnight at 4 °C. Embryos were washed with PBDT (PBS 1x, BSA 1 percent, DMSO 1 percent, and Triton X-100 0.3 percent) and blocked for 1-2 hours in blocking buffer (PBDT and 0.5 percent goat serum). Primary antibodies were diluted in blocking buffer and incubated overnight at 4 °C. Embryos were rinsed several times with PBDT and incubated with secondary antibodies for 4 h at room temperature. Embryos were rinsed in PBDT and mounted in 50 percent Glycerol/PBS. Following concentrations were used: ZO-1 (1:250), Acetylated tubulin (1:500), anti-mouse IgG2b Alexa Fluor 488 (1:500) and anti-mouse IgG1 Alexa Fluor 594 (1:500).

### Zebrafish injection

Translation blocking morpholinos (Gene Tool LLC Philomath) were injected in 1-cell stage embryos with 2nL of injection volume and 0.5 mM concentration. Following morpholinos were used: minibar MO (5’-GTATTCAATGAGCCCAAGGAGTGTC-3’); minibar MO-2 (5’-TCACCAAGCGGACACCTTTCAATGC-3’); standard control (5-CCTCTTACCTCAGTTACAATTTATA- 3’). For rescue experiments, pCS2+-Minbar-GFP plasmid was linearized with NotI and Capped MiniBAR-GFP mRNA was synthetized using mMessage mMachine SP6 kit (Thermo Fischer). MiniBAR-GFP mRNA was injected in 1-cell stage embryos with 2 nl of injection volume and 90 ng/µL.

### CRISPR/Cas13d in zebrafish embryos

Three different guide RNAs targeting minibar were prepared following Kushawah et al. protocol (Kushawah et al., 2020). Targeted sequences can be found in the following table. RfxCas13d protein, purified from bacteria cells transformed with the plasmid pET-28b-RfxCas13d-His (Addgene 141322), was kindly provided by JP Concordet. RfxCas13d purified protein and a mix of 3 guide RNAs targeting minibar were injected in 1-cell stage embryos with 2 nl of injection volume and 600 ng/µL (gRNAs) and 3µg/µL (Cas13d) concentrations. RfxCas13d purified protein was injected alone as a control.

### Statistics and reproducibility data

All values are displayed as mean ± SD (standard deviation) for at least three independent experiments (as indicated in the figure legends). Significance was calculated using paired two-sided t-tests, or one-sided exact Fisher’s tests or mixed linear model, as indicated in the figure legends. For comparing distribution, a nonparametric Kolmogorov–Smirnov (KS) test was used. P values are indicated in each individual graph and were rounded to the nearest significant figure. In all statistical tests P>0.05 was considered as non-significant. Quantifications of images and Western blots were done with Fiji.

### Data availability

The atomic model of the Rac1BD and Rab35BD are available on the PDB under the accession code 8BUY and 8BUX, respectively. The other data supporting the findings of this study are available from the corresponding author upon request.

## ACKNOWLEDGEMENTS

We thank Philippe Bastin, Renata Basto, Alexandre Benmerah and members of our Labs for critical reading, discussions and suggestions; Chantal Desdouets for antibodies, Alexandre Benmerah for plasmids and antibodies, Claire Wyart for the Tg(actb2:Mmu.Arl13b-GFP) line, Marion Delous for the Tg(wt1b:gfp) line, Christine Vesque for the spaw and pitx2 probes, Jean-Paul Concordet for the Cas13d protein, Bruno Goud and Stéphanie Miserey-Lenkei for the Yeast 2-hybrid Rab GT-Pase library. We thank Anne-Marie Wehenkel and Daniela Megrian Nunez for help with AlphaFold; Chiara Zurzolo and Inés Saenz-de-Santa-Maria for commissioning the moving kymograph tool, Pierre Lafaye for initial immunization experiments, Pierre-Henri Commere from Cytometry and Biomarkers UTechS, Institut Pasteur for FACS sorting, Willy Supatto for the help with cilia motility imaging, Emilie Menant for fish care, P. Mahou and the Polytechnique Bioimaging Facility for imaging on their equipment supported by Région Ile-de-France (interDIM) and Agence Nationale de la Recherche (ANR-11-EQPX-0029 Morphoscope2, ANR-10-INBS-04 France BioImaging). UTechS PBI is part of the France–BioImaging infrastructure network (FBI) supported by the French National Research Agency (ANR-10-INBS- 04; Investments for the Future), and acknowledges support from ANR/FBI and the Région Ile-de-France (program “Domaine d’Intérêt Majeur-Malinf”) for the use of the Zeiss LSM 780 Elyra PS1 microscope. We acknowledge SOLEIL for provision of synchrotron radiation facilities, and we would like to thank Pierre Legrand for assistance in using the "Proxima-1" beamline and for outstanding help in data processing and structure determination; William Shepard, Martin Savko, Serena Sirigu for assistance in using the "Proxima-2" beamline; Aurelien Thureau for assistance in using the "SWING" beamline and approval of data analysis. We thank Carlos Kikuti and Cecile Boutonnet for help and support in the SEC-MALS experiments, the IBENS platform for access to MicroCal-iTC200 instrument and David Stroebel for assistance in ITC data collection and processing. This work has been supported by the Institut Pasteur, the CNRS, ANR 18-CE13-0024, ANR 19-CE13- 0018 and Fondation pour le Recherche Médicale: Recherche soutenue par la FRM EQU202103012627 to AE, ANR 18-CE13-0024 and ANR 20-CE13-0016 to ND, ANR 18-CE13-0024, INCa 2014-1-PL BIO-04-ICR-1, FRM ING20140129255 to AH and ANR-20-CE18-0016-02 to OP. The Structural Motility Team is part of the Labex «Cell(n)Scale» with the references ANR-10-LABX-0038 and ANR-10-IDEX-0001-02. HH has been awarded a doctoral fellowship from the PSL Université. SE was supported by the European Union’s Horizon 2020 programme under the Marie Skłodowska-Curie grant agreement No 840201. RS received a fellowship from Association de Recherche sur le Cancer ARC PDF20171206712.

---

**Figure S1:**
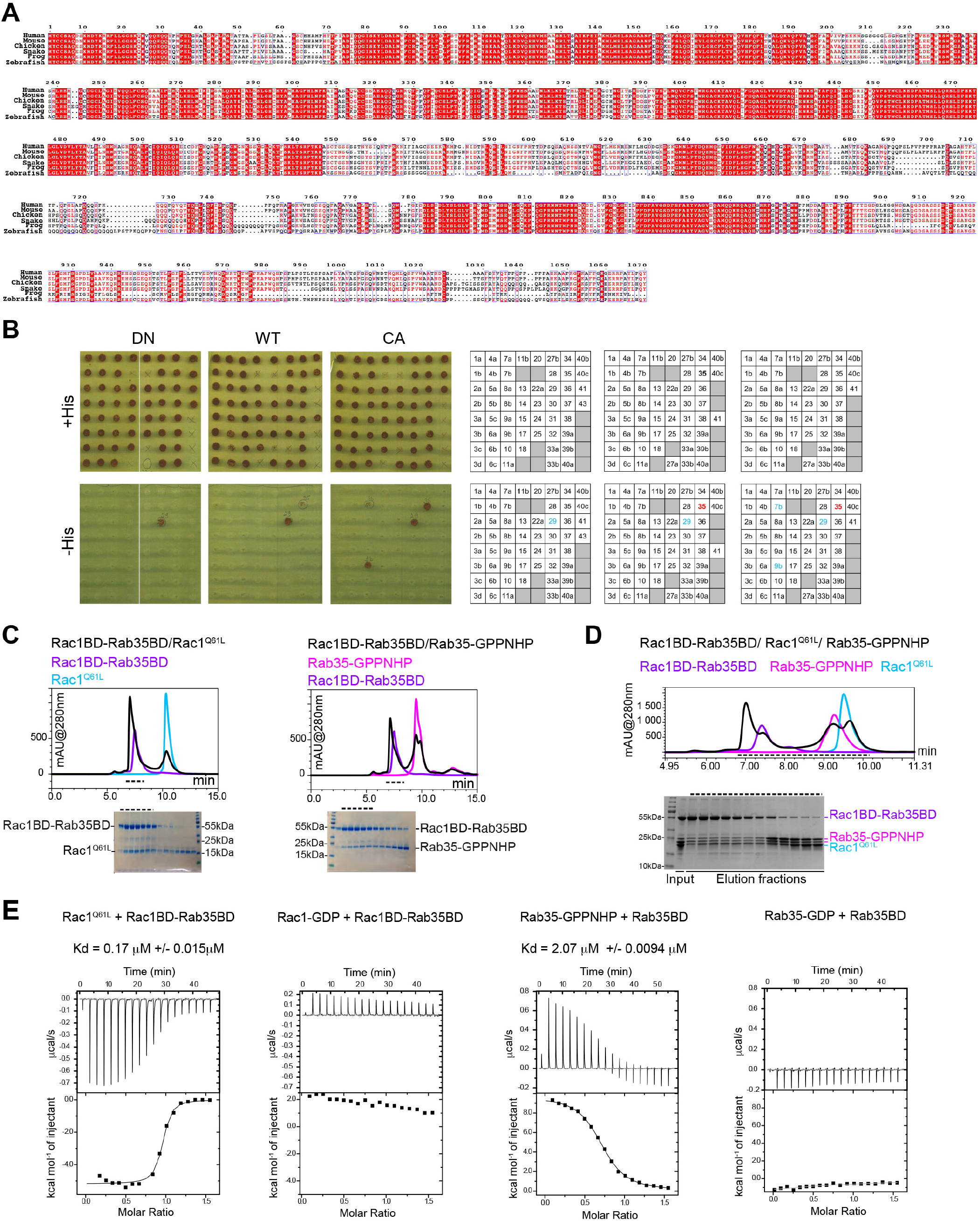
Alignment of full-length MiniBAR proteins from different species. Specificity of MiniBAR interaction with Rab35. Biochemical characterization of Rac1 and Rab35 interaction with MiniBAR domains. (Related to Figure 1) (**A**) The N-terminal part of the protein (MiniBAR70-553) that binds both Rac1 and Rab35 GTPases is highly conserved in vertebrates, showing 76 percent identity and 85.5 percent similarity between human and fish sequences. The C-terminal part of the protein (MiniBAR554-1070) is predicted to be intrinsically disordered.UniProt accession numbers: Human (Homo sapiens): O15063; Mouse (Mus musculus) Q6PAL5, Chicken (Gallus gallus): E1BT43; Frog (Xenopus tropicalis): A0A6I8S090; Snake (Pseudonaja textilis): A0A670YC05 and Zebrafish (Danio rerio): A0A0R4IGD5. (**B**) S. cerevisiae L40 reporter strain was transformed with plasmids encoding Gal4 Activation Domain (GAD) fused to MiniBAR Rab35BD to examine interactions with LexA fused to 55 different human Rab GTPases either wild type (WT) or dominant negative (DN, mutation equivalent to S22N in Rab35) and constitutively active (CA, mutation equivalent to Q67L in Rab35). Growth on a medium without histidine (- His) indicates an interaction with the corresponding proteins in this two-hybrid assay. Among all tested Rab GTPase, only Rab35 WT and CA showed a specific growth (red numbers). Growth with Rab29 and Rab9b is not conclusive since it is observed with any GAD fusions (constitutive growth due to autoactivation by Rab29 and Rab9, blue numbers). (**C**) Analytical gel-filtration elution profiles of individual proteins: Rac1BD-Rab35BD fragment (MiniBAR70-553), Rab35- GPPNHP and Rac1Q61L in comparison with their complexes. The elution fractions corresponding to the complexes elution peaks were analyzed by SDS-PAGE (marked with black dashed lines), masses of the eluted proteins and complexes were determined by MALS and indicated below. Left panel: Rac1BD-Rab35BD (MassMALS 97.2 ± 0.7 percent kDa), Rac1Q61L (MassMALS 20.6 ± 0.3 percent kDa), Rac1BD-Rab35BD/Rac1Q61L complex (MassMALS 123.8 ± 0.4 percent kDa). Left panel: Rac1BD-Rab35BD (MassMALS 97.2 ± 0.7 percent kDa), Rab35-GPPNHP (MassMALS 22.2 ± 0.1 percent kDa), Rac1BD-Rab35BD/Rab35-GPPNHP complex (MassMALS 117.1 ± 0.5 percent kDa). (**D**) Analytical gel-filtration elution profiles of individual proteins Rac1BD-Rab35BD, His-tagged-Rab35-GPPNHP and Rac1Q61L in comparison with their complexes. His-tagged-Rab35 migrates as a double band. Both GTPases co-elute with the Rac1BD-Rab35BD fragment. (**E**) The ITC curves show the interactions between the active or inactive states of Rac1 with Rac1BD-Rab35BD (left panels) and the active or inactive states of Rab35 and Rab35BD (right panels)

**Figure S2:**
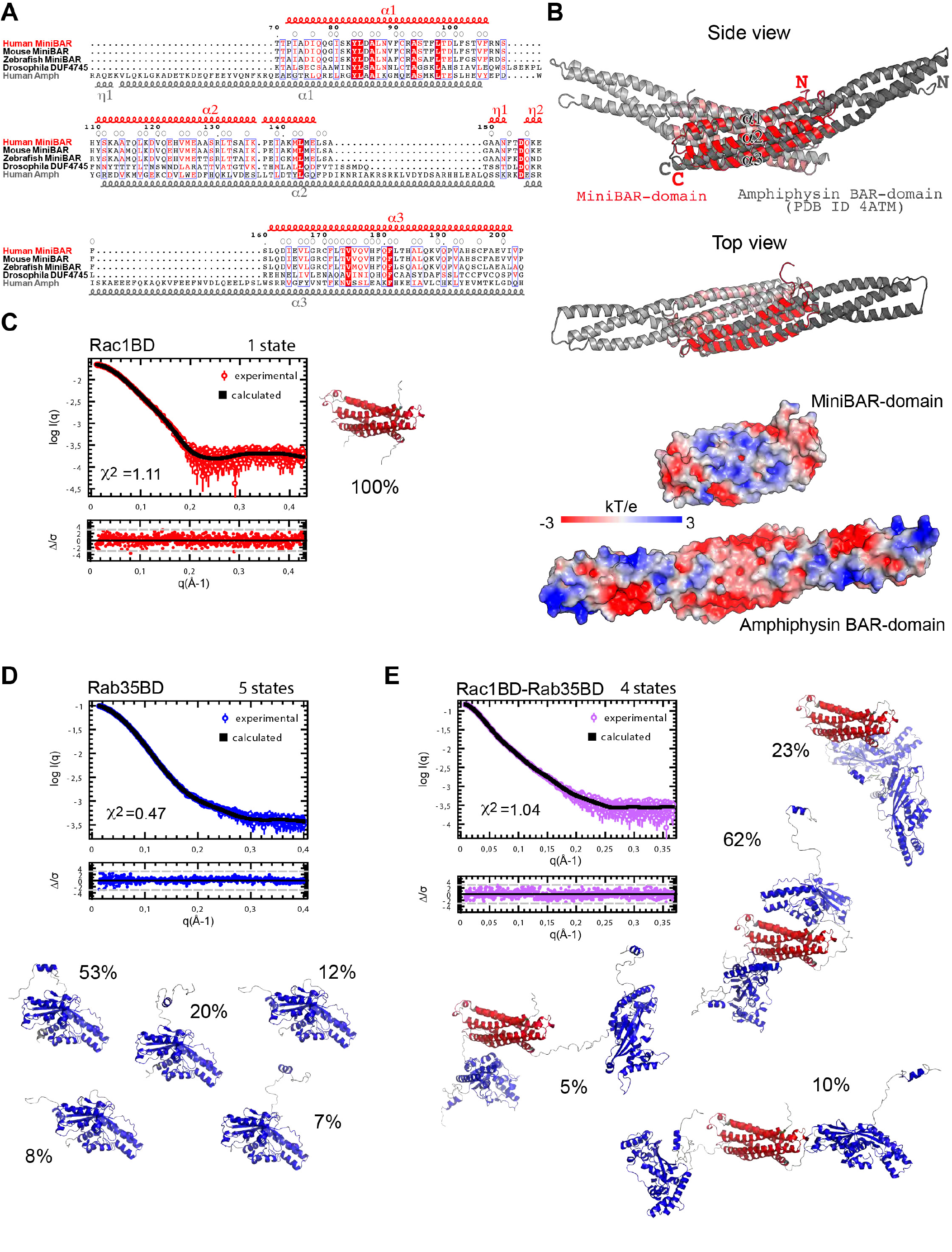
Comparison of the truncated BAR domain of MiniBAR and the BAR domain of Amphiphysin. Models of MiniBAR fragments fitted to the SAXS data. (Related to Figure 1) (**A**) Sequence alignment of the truncated BAR domains from human, mouse and fish MiniBAR with DUF4745 domain (http://pfam.xfam.org/family/DUF4745) from D. melanogaster protein (B5RJJ4) and human Amphyphisin BAR domain. Residues participating to the homodimer formation are marked with open circles. Secondary structure elements distribution of the Human MiniBAR and Human Amphiphysin are shown in red above the alignment and in grey below the alignment, respectively. (**B**) Top panels: Side view and Top view of the superimposition of the truncated BAR domain of human MiniBAR (red) on the core part of the canonical human Amphiphysin BAR domain (in grey), a concave homodimer made of 2×3 alpha-helices usually present in proteins that sense and/or induce membrane curvature. The MiniBAR homodimer interface is mostly hydrophobic. MiniBAR helix-2 and helix-3 are kinked at the positions of Pro-137 and Pro-191 respectively helping the typical BAR domain monomers mutual wrapping. The MiniBAR connection loops between helix-1 and helix-2 are short, similar to the Amphiphysin structure; in contrast, the helix-2 and helix-3 are connected with a longer flexible loop, replacing the “arm” regions formed by the extensions of the helix-2 (C-terminal part), helix-3 (N-terminal part) and helix-1 (N-terminal part) of Amphiphysin. Bottom panels: Electrostatic potential distribution maps on the surfaces of truncated BAR of human MiniBAR and the canonical BAR of human Amphiphysin at the top view orientation. (**C-E**) Models of MiniBAR fragments fitted to the SAXS data using MultiFoXS algorithm. Fitting curves and error weighted residual plots of the models. Different states representative models and the state populations are also shown: 1 state for Rac1BD (C), 5 states for Rab35BD (D) and 4 states for Rac1BD-Rab35BD (E).

**Figure S3:**
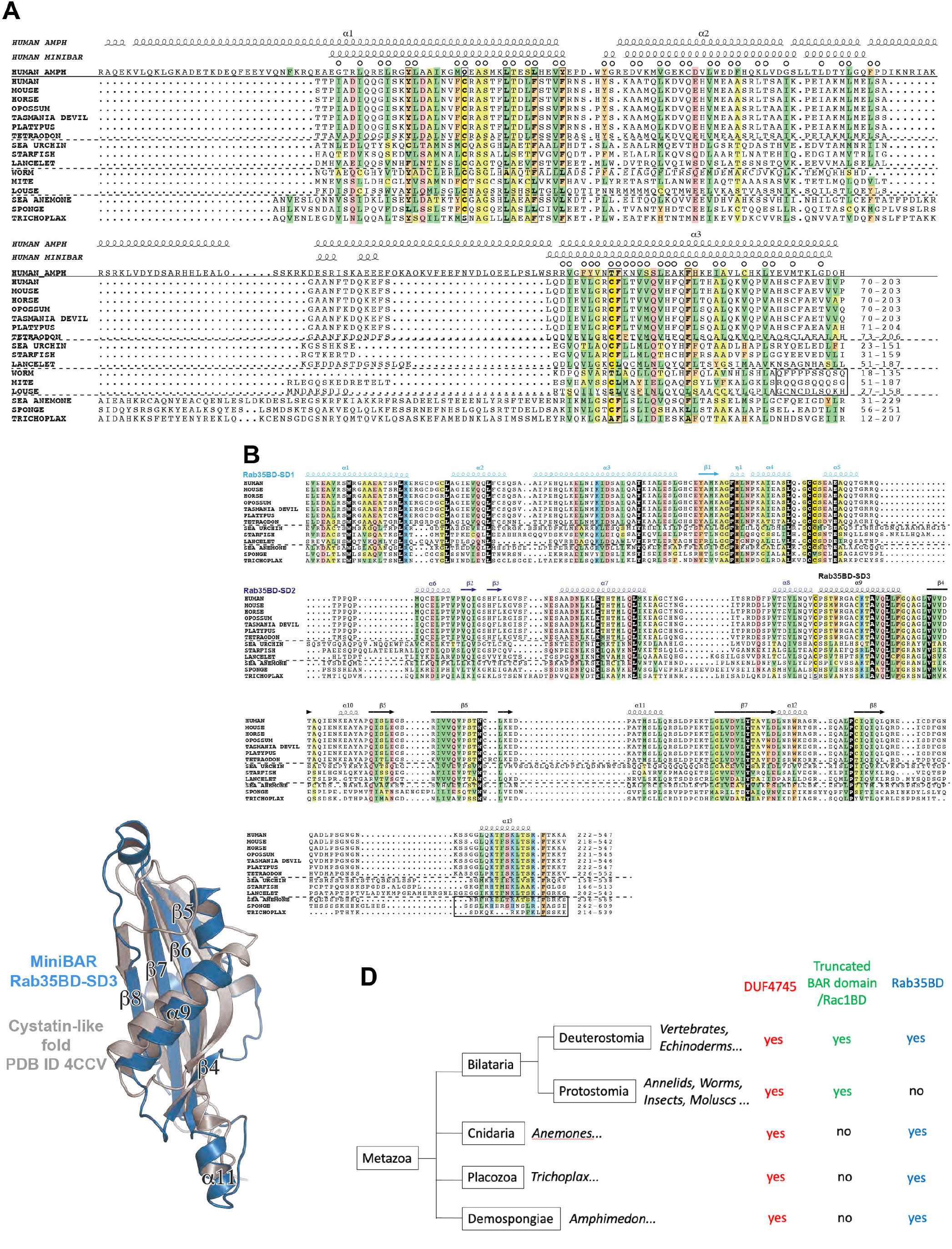
Alignment of full-length MiniBAR, Rac1BD and Rab35BD in multiple species across evolution. Comparison of Rab35BD-SD3 and a cystatin-like fold. (Related to Figure 1) (**A-B**) Sequence alignments of Rac1BD (A) and Rab35BD (B) from different species, based on MAFFT run on the EMBL-EBI server(Madeira et al., 2022) and rendered using ESPript(Robert and Gouet, 2014). In (A), the sequence of Amphiphysin, which displays a normal BAR domain, is given for comparison. Note that Sponges, Placozoa and Cnidaria have a normal BAR domain. The secondary structures of experimental 3D structures (Human Amphiphysin: pdb 4ATM, Human MiniBAR: this study) are reported above the alignment, with residues participating in the dimer formation indicated with open circles, as in Figure S2A. Boxes indicate more variable regions, with uncertain alignment and for which differences are observed in the AF2 models. Colors are used to highlight positions for which conserved features observed (green: strong hydrophobic amino acids (V, I, L, M, F, Y, W); orange: aromatic amino acids (F, Y, W, H); light green: small, non-strong hydrophobic amino acids (A,G,C) and amino acids that can substitute for them (S, T); red : D,E,Q,N,S,T; blue : K,R,H). UniProt accession numbers: HUMAN (Homo sapiens): O15063; MOUSE (Mus musculus): Q6PAL5; HORSE (Equus caballus): F7DL01; OPOSSUM (Monodelphis domestica): F7B911; TASMANIA DEVIL (Sarcophilus harrisii): A0A7N4NPN4; PLATYPUS (Ornithorhynchus anatinus): F7BGA3; TETRAODON (Tetraodon nigroviridis): H3CYU6; SEA URCHIN (Strongylocentrotus purpuratus): A0A7M7N6V8; STARFISH (Acanthaster planci): A0A8B7YHJ9; LANCELET (Branchiostoma belcheri): A0A6P5A9Y3; WORM (Capitella teleta): R7TMM2; MITE (Varroa destructor): A0A7M7J8J6; LOUSE (Pediculus humanus): E0W1T5; SEA ANEMONE (Actinia tenebrosa): A0A6P8H0T2; SPONGE (Amphimedon queenslandia): A0A1X7V5V2; TRICHOPLAX (Trichoplax adhaerens): B3RJE9. (**C**) Rab35BD SD-3 superimposed on a cystatin fold representative structure (PDB 4CCV) using vector alignment search tool(Madej et al., 2020). In Rab35BD, the N-terminal SD-1 is mostly alpha-helical, it contains 5 alpha helices (alpha1-alpha5) and only one beta strand (beta1). SD-2 is formed by 3 helices (alpha6-alpha8) and two antiparallel beta strands (beta2 and beta3). In Rab35BD, the C-terminal subdomain (SD-3) is formed by an extended 4-stranded beta sheet with two alpha helices (alpha9 and alpha12) on the opposite surfaces of the sheet and two helical inserts (alpha10 and alpha11) in the loops connecting the beta strands. Beta-strand from SD-1 extends the central SD-3 beta-sheet at one edge and the two SD-2 beta strands from another edge, thus forming a single 7 stranded mixed beta sheet. (**D**) Presence of DUF4745, of a truncated BAR domain/Rac1BD and of a Rab35BD across evolution. Note that the DUF4745 is ancient in metazoa, and that the truncated BAR domain is specific of Bilateria. The Rab35BD is ancient and has been lost in Protostomia. Thus, only Deuterostomia (which include Echinoderms, Cephalochordates, Tunicates and Vertebrates) have a truncated BAR domain followed by a Rab35BD.

**Figure S4:**
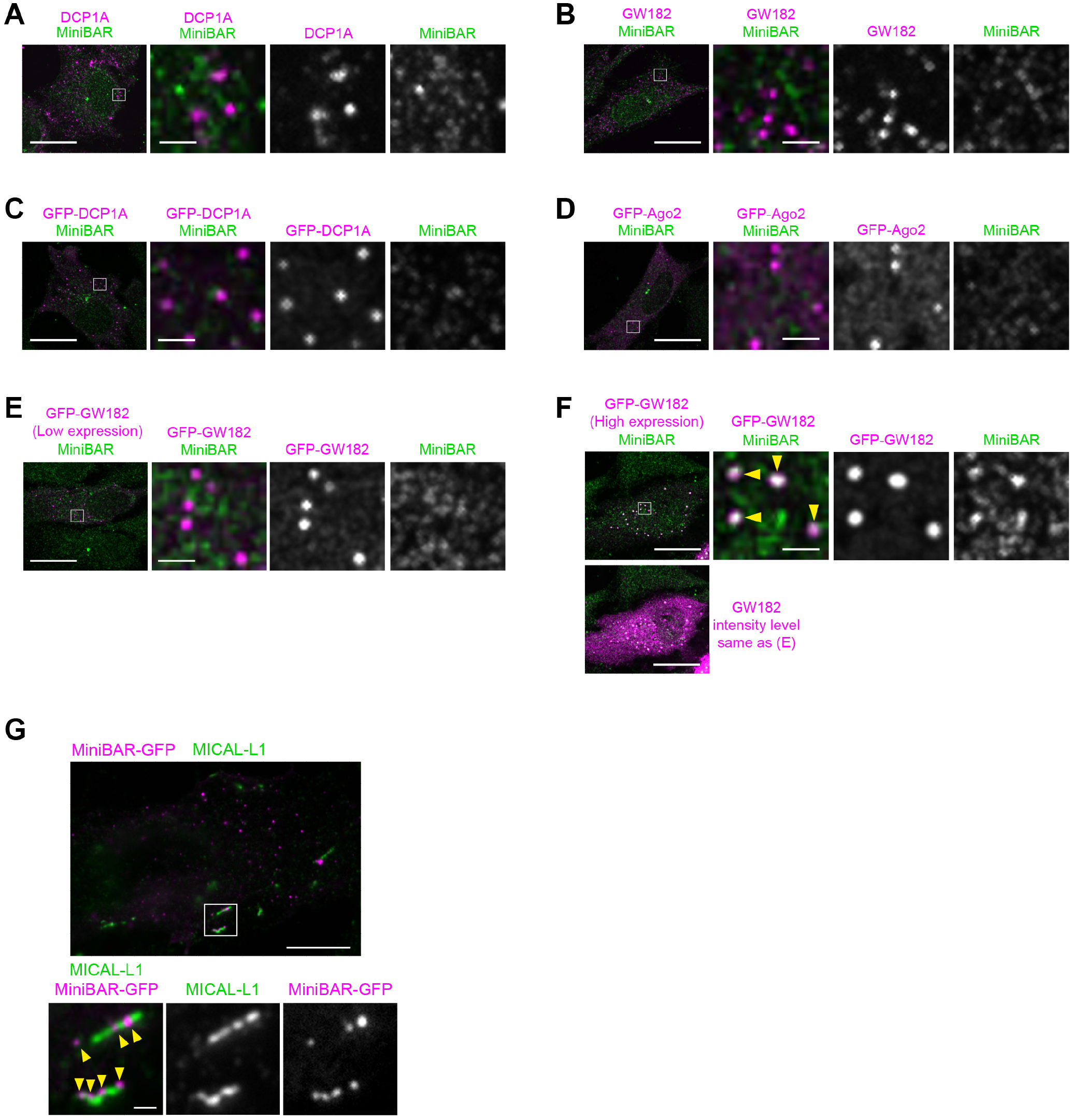
Respective localization of MiniBAR and P-granule markers. Localization of MiniBAR and MICAL-L1. (Related to Figure 2) (**A**) Representative image of RPE-1 cells stained with endogenous MiniBAR and endogenous DCP1A. Scale bars, 20 µm (general view) and 4 µm (insets of the boxed regions). (**B**) Representative image of RPE-1 cells stained with endogenous MiniBAR and endogenous GW182. Scale bars, 20 µm (general view) and 4 µm (insets of the boxed regions). (**C**) Representative image of RPE-1 cells transfected with plasmids encoding GFP-DCP1A and stained for endogenous MiniBAR. Scale bars, 20 µm (general view) and 4 µm (insets of the boxed regions). (**D**) Representative image of RPE-1 cells transfected with plasmids encoding GFP-Ago2 and stained for endogenous MiniBAR. Scale bars, 20 µm (general view) and 4 µm (insets of the boxed regions). (**E-F**) Representative images of RPE-1 cells transfected with plasmids encoding GFP-GW182 at low levels (E) or high levels (**F**) and stained for endogenous MiniBAR. Scale bars, 20 µm (general view) and 4 µm (insets of the boxed regions). In (F) top panels, the display of the magenta channel has been reduced for better visualization of the colocalization with MiniBAR. The magenta levels in the bottom picture have been set as in (E) for comparison of the expression levels. In A-F, note that endogenous MiniBAR did not co-localize with endogenous markers of P-granules (DCP1A, GW182) (A-B) nor with GFP-DCP1A, GFP-Ago2 or GFP-GW182 when expressed at low levels (C-E). Only when over-expressed at high levels, did GFP-GW182 (but not GFP-DCP1 or GFP-Ago2) colocalize with endogenous MiniBAR (F). (**G**) Representative image of RPE-1 cells transfected with plasmids encoding MiniBAR-GFP and stained for endogenous MICAL-L1, as indicated. Scale bars, 20 µm (general view) and 2 µm (insets of the boxed regions).

**Figure S5:**
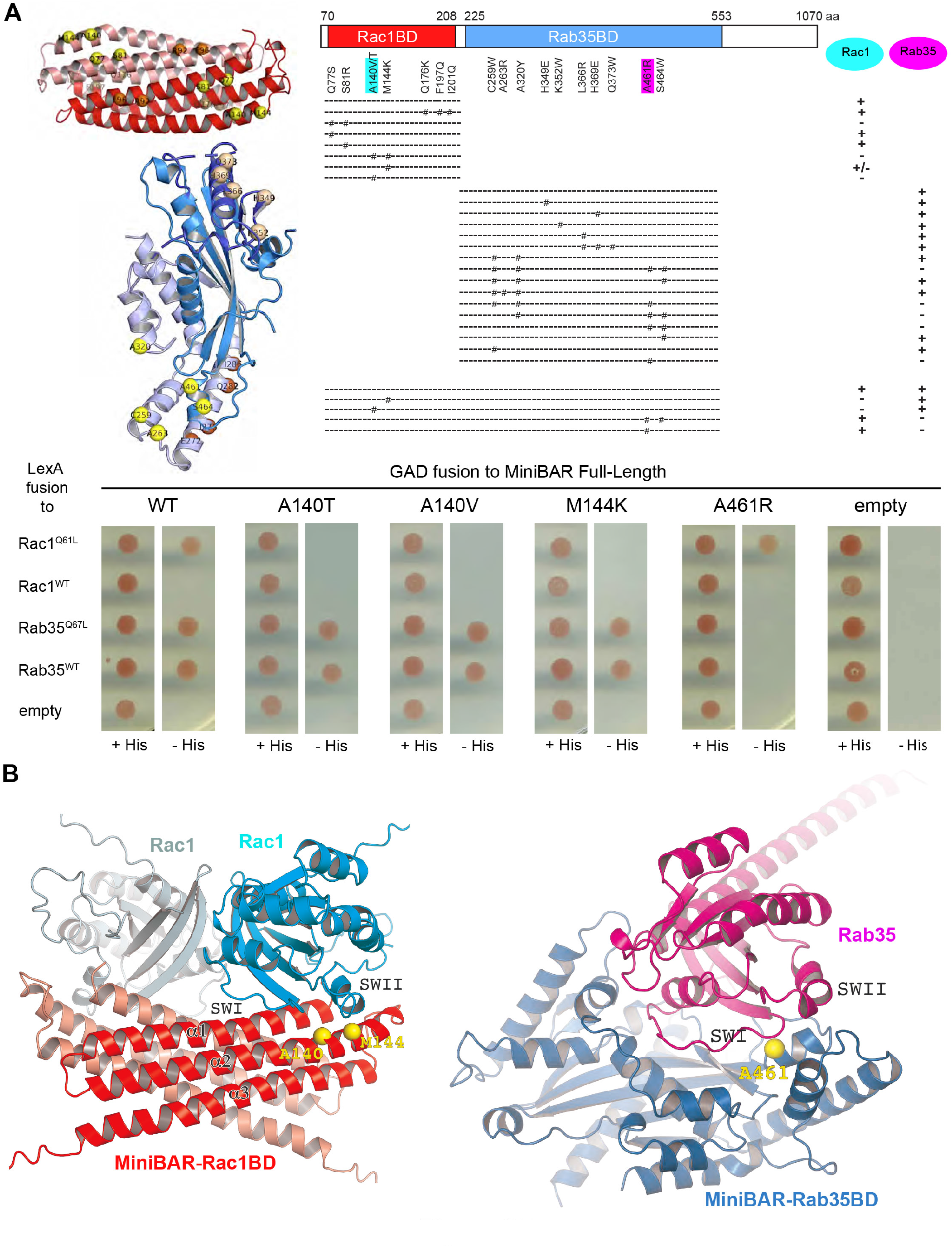
Screening for point mutations that selectively disrupt MiniBAR interactions with either Rac1 or Rab35. (Related to Figure 2) (**A**) Left panel: The localization of the aa mutated in the Rac1BD (top) and in the Rab35BD (bottom) are shown as spheres. Right panel: Summary of the combined and single aa substitutions in the Rac1BD, the Rab35BD and in the full-length MiniBAR tested in yeast 2-hybrid for interactions with either LexA-Rac1Q61L or LexA-Rab35 Q67L fusions. +, ± and - indicate normal, weak and no growth of L40 reporter strain in the absence of Histidine, respectively. Bottom panel: results of the growth of the reporter L40 strain of single amino acid substitution in full-length MiniBAR, as indicated. Growth on a medium without histidine (- His) indicates an interaction with the corresponding proteins in this two-hybrid assay. (**B**) Left panel: AlphaFold predicted complex structure of Rac1BD with Rac1. The prediction is consistent with the A140 and M144 involvement in Rac1 binding shown in (A) and Figure 2E. Right panel: AlphaFold predicted complex structure of Rab35BD with Rab35. The prediction is consistent with the A461 involvement in Rab35 binding shown in (A) and Figure 2F.

**Figure S6:**
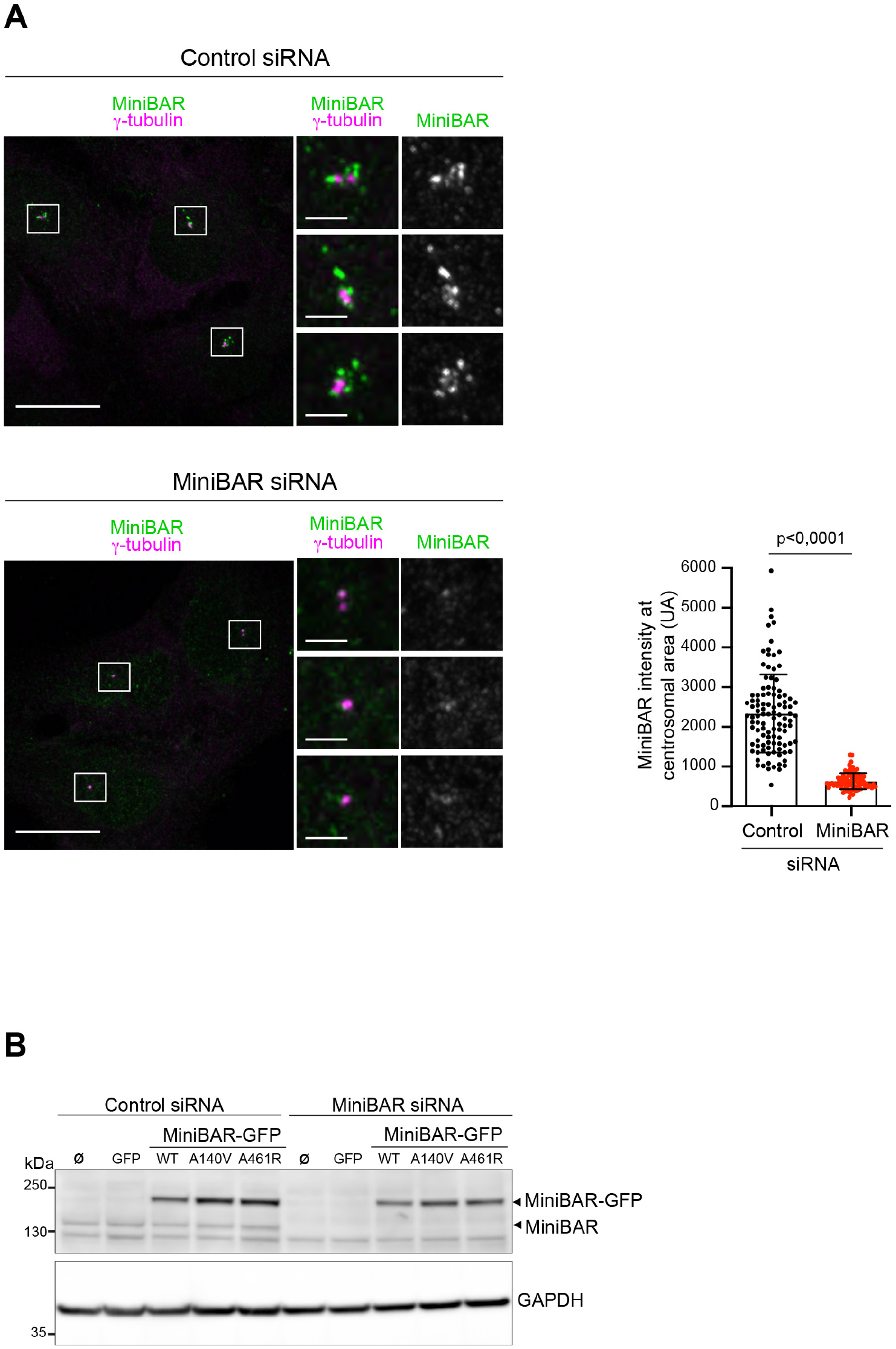
Specificity of MiniBAR staining in RPE cells. Levels of MiniBAR in stably expressing cells. (Related to Figure 3) (**A**) Representative images of RPE-1 cells transfected with either control or MiniBAR siRNAs and stained for endogenous MiniBAR and G-Tubulin. Zoomed regions of the centrosomal area are displayed as well as the quantification of the mean intensity of MiniBAR at the centrosomal area. Scale bars, 20 µm (general view) and 2 µm (insets of the boxed regions). (**B**) Lysates of RPE-1 cells stably expressing the indicated constructs and transfected with either control or MiniBAR siRNAs were blotted for MiniBAR and GAPDH (loading control).

**Figure S7:**
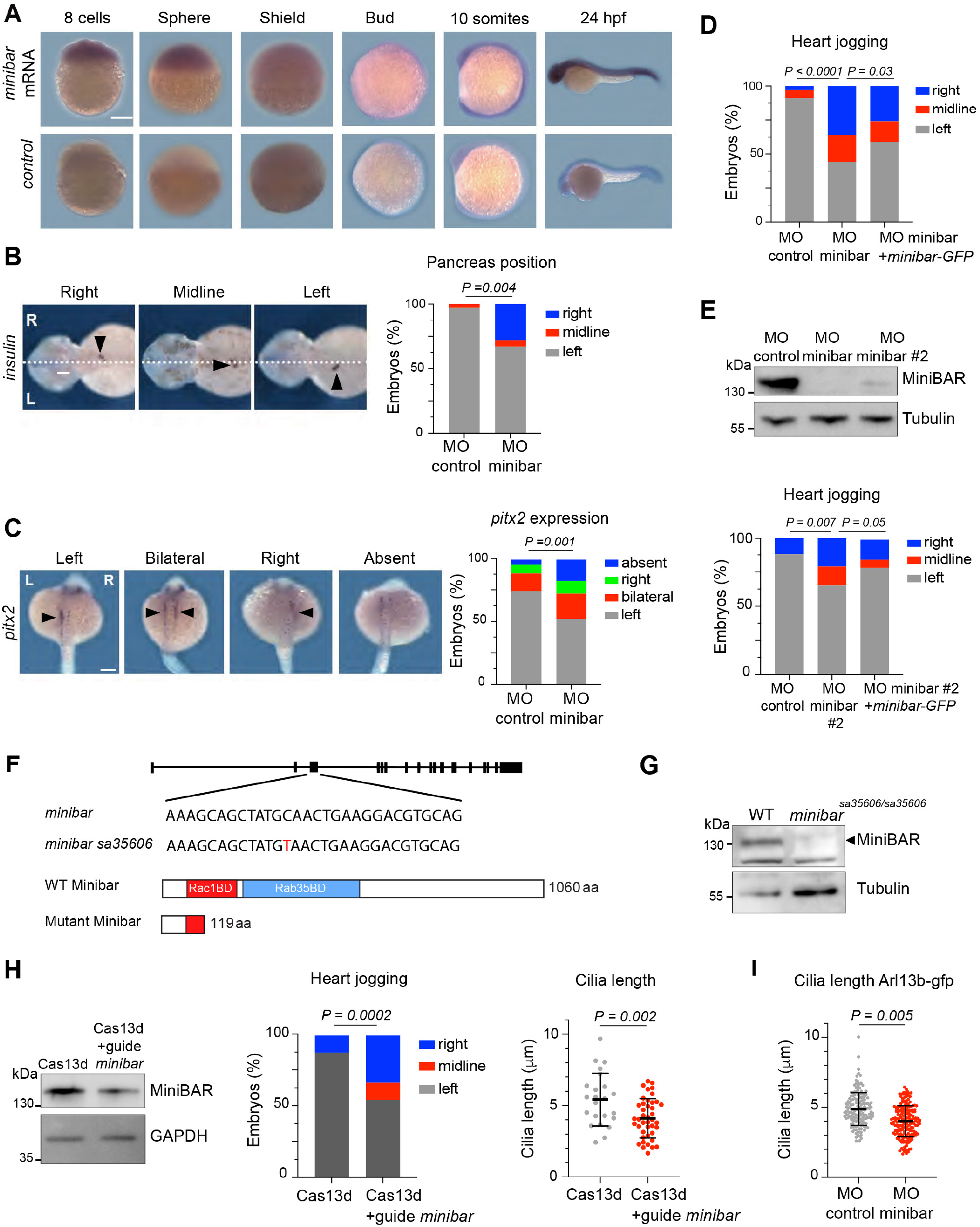
Expression of minibar mRNA during early development. L/R asymmetry defects observed after MiniBAR depletion using several approaches. (Related to Figure 6) (**A**) Expression of minibar visualized by in situ hybridization in zebrafish embryos from the 8-cell stage to 24 hpf. Control is the sense probe. Scale bars, 200 µm. (**B**) In situ hybridization of the pancreatic specific mRNA markers insulin at 48 hpf. Scale bar, 100 µm. Quantification of left, midline and right expression of insulin. n = 33 and 21 embryos for control and minibar MO, respectively. Chi-square test. (**C**) pitx2 expression at the 23-somite stage. Scale bar, 200 µm. Quantification of left, bilateral, right and absent expression of pitx2. n = 59 and 34 embryos for control and minibar MO, respectively. Fischer exact test. (**D**) Co-injection of minibar-GFP mRNAs with the minibar morpholino partially rescues the heart jogging phenotype. n = 68 embryos for control MO, 142 for minibar MO and 135 for minibar MO with minibar-GFP mRNAs, respectively. Chi-square test. The histogram in Fig. 6e is a truncated version of this histogram. (**E**) Left panel: Lysates of 24 hpf zebrafish embryos injected with either a control morpholino (MO control), the morpholino targeting minibar (MO minibar) used in Figure 6 or a second minibar morpholino (MO minibar 2) were blotted for MiniBAR and Tubulin (loading control). The blot in Figure 6A is a truncated version of this blot. Exposure time has been increased here to show the differences in MiniBAR depletion obtained with the two MOs. Right panel: effect of MO minibar 2 on heart jogging, and partial rescue by co-injection of minibar-GFP mRNAs. n = 43 embryos for control MO, 114 for minibar MO2 and 124 for minibar MO2 with minibar-GFP mRNAs. Chi-square test. (**F**) minibarsa35606 is a point mutation in the third exon, inducing a premature stop codon in the Rac1 binding domain. (**G**) Lysates of 24 hpf wild type (WT) or minibarsa35606/sa35606 embryos were blotted for MiniBAR and Tubulin (loading control). (**H**) Injection of Cas13 with guide RNAs targeting minibar leads to a reduction of MiniBAR expression (Western blot, left panel), heart jogging defects (middle panel) and a reduction of cilia length in Kupffer’s vesicle (right panel). (**I**) Cilia length in the Kupffer’s vesicle in -actin2:arl13b-gfp embryos injected with either control or minibar MOs. n = 9 embryos and 159 cilia for control and 9 embryos and 187 cilia for minibar MO.

**Table S1:**
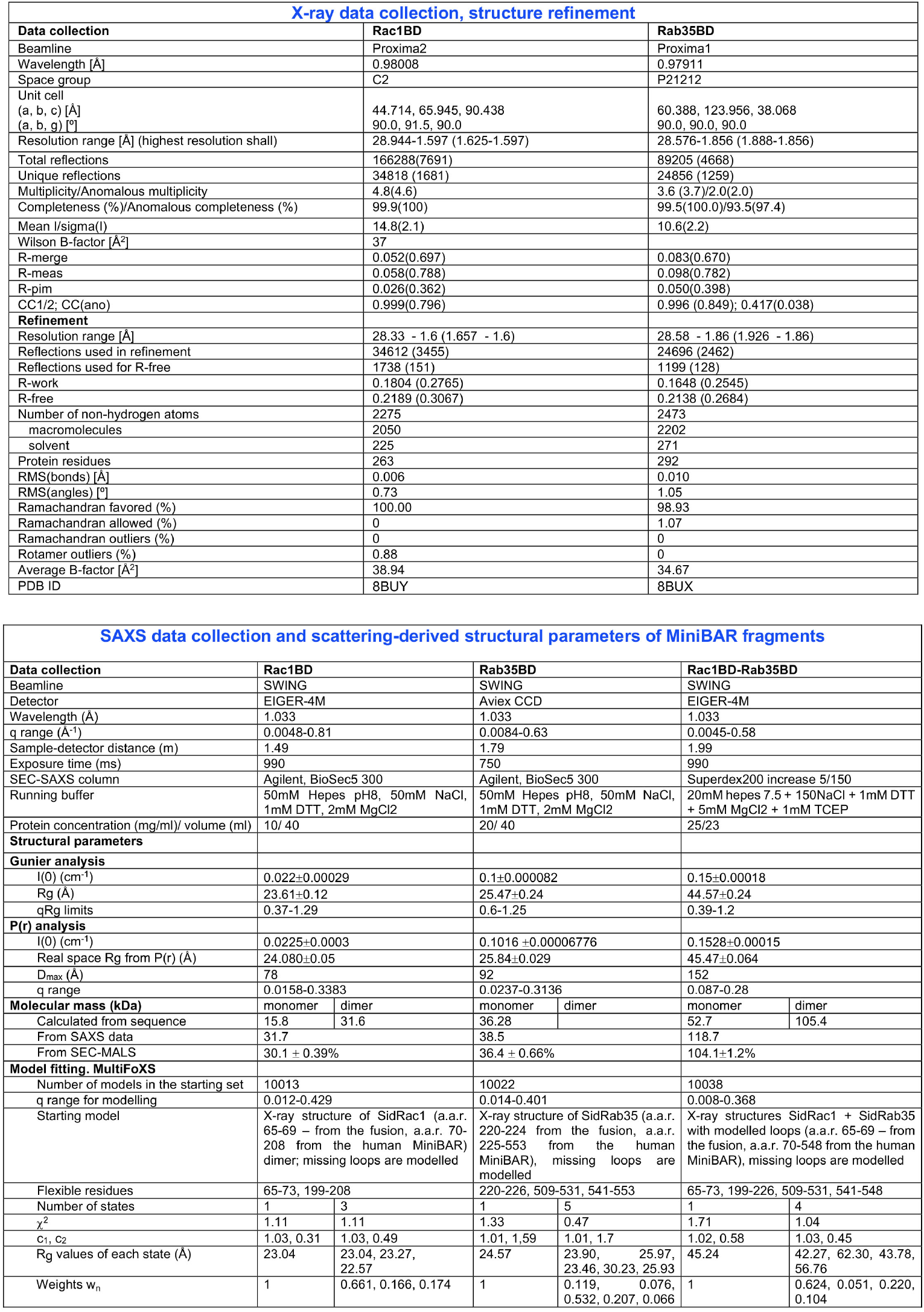
X-ray and SAXS data collection (related to Figure 1)

## References

Adams, P.D., Afonine, P.V., Bunkóczi, G., Chen, V.B., Davis, I.W., Echols, N., Headd, J.J., Hung, L.W., Kapral, G.J., Grosse-Kunstleve, R.W., et al. (2010). PHENIX: a comprehensive Python-based system for macromolecular structure solution. Acta Crystallogr D Biol Crystallogr 66, 213–221. 10.1107/s0907444909052925.

Allaire, P.D., Marat, A.L., Dall’Armi, C., Di Paolo, G., McPherson, P.S., and Ritter, B. (2010). The Connecdenn DENN domain: a GEF for Rab35 mediating cargo-specific exit from early endosomes. Mol Cell 37, 370–382. S1097–2765(10)00076-6 [pii] 10.1016/j.molcel.2009.12.037.

Allaire, P.D., Seyed Sadr, M., Chaineau, M., Seyed Sadr, E., Konefal, S., Fotouhi, M., Maret, D., Ritter, B., Del Maestro, R.F., and McPherson, P.S. (2013). Interplay between Rab35 and Arf6 controls cargo recycling to coordinate cell adhesion and migration. J Cell Sci 126, 722–731. 10.1242/jcs.112375.

Anvarian, Z., Mykytyn, K., Mukhopadhyay, S., Pedersen, L.B., and Christensen, S.T. (2019). Cellular signalling by primary cilia in development, organ function and disease. Nature reviews. Nephrology 15, 199–219. 10.1038/s41581-019-0116-9.

Bagci, H., Sriskandarajah, N., Robert, A., Boulais, J., Elkholi, I.E., Tran, V., Lin, Z.Y., Thibault, M.P., Dubé, N., Faubert, D., et al. (2020). Mapping the proximity interaction network of the Rho-family GTPases reveals signalling pathways and regulatory mechanisms. Nature cell biology 22, 120–134. 10.1038/s41556-019-0438-7.

Ball, G., Demmerle, J., Kaufmann, R., Davis, I., Dobbie, I.M., and Schermelleh, L. (2015). SIMcheck: a Toolbox for Successful Super-resolution Structured Illumination Microscopy. Scientific reports 5, 15915. 10.1038/srep15915.

Barral, D.C., Garg, S., Casalou, C., Watts, G.F., Sandoval, J.L., Ramalho, J.S., Hsu, V.W., and Brenner, M.B. (2012). Arl13b regulates endocytic recycling traffic. Proceedings of the National Academy of Sciences of the United States of America 109, 21354–21359. 10.1073/pnas.1218272110.

Bergfors, T. (2003). Seeds to crystals. J Struct Biol 142, 66–76. 10.1016/s1047-8477(03)00039-x.

Blacque, O.E., Scheidel, N., and Kuhns, S. (2018). Rab GT-Pases in cilium formation and function. Small GTPases 9, 76–94. 10.1080/21541248.2017.1353847.

Bollig, F., Perner, B., Besenbeck, B., Köthe, S., Ebert, C., Taudien, S., and Englert, C. (2009). A highly conserved retinoic acid responsive element controls wt1a expression in the zebrafish pronephros. Development 136, 2883–2892. 10.1242/dev.031773.

Borovina, A., Superina, S., Voskas, D., and Ciruna, B. (2010). Vangl2 directs the posterior tilting and asymmetric localization of motile primary cilia. Nature cell biology 12, 407–412. 10.1038/ncb2042.

Breslow, D.K., Koslover, E.F., Seydel, F., Spakowitz, A.J., and Nachury, M.V. (2013). An in vitro assay for entry into cilia reveals unique properties of the soluble diffusion barrier. The Journal of cell biology 203, 129–147. 10.1083/jcb.201212024.

Bricogne G., B.E., Brandl M., Flensburg C., Keller P., Pa-ciorek W.„ and Roversi P, S.A., Smart O.S., Vonrhein C., Womack T.O. (2017). BUSTER version 2.10.3 Cambridge,. United Kingdom: Global Phasing Ltd.

Cai, J., Song, X., Wang, W., Watnick, T., Pei, Y., Qian, F., and Pan, D. (2018). A RhoA-YAP-c-Myc signaling axis promotes the development of polycystic kidney disease. Genes Dev 32, 781–793. 10.1101/gad.315127.118.

Carman, P.J., and Dominguez, R. (2018). BAR domain proteins-a linkage between cellular membranes, signaling pathways, and the actin cytoskeleton. Biophys Rev 10, 1587–1604. 10.1007/s12551-018-0467-7.

Caspary, T., Larkins, C.E., and Anderson, K.V. (2007). The graded response to Sonic Hedgehog depends on cilia architecture. Developmental cell 12, 767–778. 10.1016/j.devcel.2007.03.004.

Cauvin, C., Rosendale, M., Gupta-Rossi, N., Rocancourt, M., Larraufie, P., Salomon, R., Perrais, D., and Echard, A. (2016). Rab35 GTPase Triggers Switch-like Recruitment of the Lowe Syndrome Lipid Phosphatase OCRL on Newborn Endosomes. Current biology : CB 26, 120–128. 10.1016/j.cub.2015.11.040.

Cevik, S., Sanders, A.A., Van Wijk, E., Boldt, K., Clarke, L., van Reeuwijk, J., Hori, Y., Horn, N., Hetterschijt, L., Wdowicz, A., et al. (2013). Active transport and diffusion barriers restrict Joubert Syndrome-associated ARL13B/ARL-13 to an Inv-like ciliary membrane subdomain. PLoS genetics 9, e1003977. 10.1371/journal.pgen.1003977.

Chaineau, M., Ioannou, M.S., and McPherson, P.S. (2013). Rab35: GEFs, GAPs and Effectors. Traffic 14, 1109–1117. 10.1111/tra.12096.

Chesneau, L., Dambournet, D., Machicoane, M., Kouranti, I., Fukuda, M., Goud, B., and Echard, A. (2012). An ARF6/Rab35 GTPase cascade for endocytic recycling and successful cytokinesis. Current biology : CB 22, 147–153. S0960-9822(11)01378-9 [pii] 10.1016/j.cub.2011.11.058.

Cowtan, K. (2000). General quadratic functions in real and reciprocal space and their application to likelihood phasing. Acta Crystallogr D Biol Crystallogr 56, 1612–1621. 10.1107/s0907444900013263.

Cowtan, K. (2006). The Buccaneer software for automated model building. 1. Tracing protein chains. Acta Crystallogr D Biol Crystallogr 62, 1002–1011. 10.1107/s0907444906022116.

Edwards, P. (2002). Origin 7.0:Scientific Graphing and Data Analysis Software. J. Chem. Inf. Comput. Sci., 1270–1271.

El-Brolosy, M.A., Kontarakis, Z., Rossi, A., Kuenne, C., Günther, S., Fukuda, N., Kikhi, K., Boezio, G.L.M., Takacs, C.M., Lai, S.L., et al. (2019). Genetic compensation triggered by mutant mRNA degradation. Nature 568, 193–197. 10.1038/s41586-019-1064-z.

Emsley, P., and Cowtan, K. (2004). Coot: model-building tools for molecular graphics. Acta Crystallogr D Biol Crystallogr 60, 2126–2132. 10.1107/s0907444904019158.

Epting, D., Slanchev, K., Boehlke, C., Hoff, S., Loges, N.T., Yasunaga, T., Indorf, L., Nestel, S., Lienkamp, S.S., Omran, H., et al. (2015). The Rac1 regulator ELMO controls basal body migration and docking in multiciliated cells through interaction with Ezrin. Development 142, 174–184. 10.1242/dev.112250.

Ershov, D., Phan, M.S., Pylvänäinen, J.W., Rigaud, S.U., Le Blanc, L., Charles-Orszag, A., Conway, J.R.W., Laine, R.F., Roy, N.H., Bonazzi, D., et al. (2022). TrackMate 7: integrating state-of-the-art segmentation algorithms into tracking pipelines. Nat Methods 19, 829–832. 10.1038/s41592-022-01507-1.

Evans, P. (2006). Scaling and assessment of data quality. Acta Crystallogr D Biol Crystallogr 62, 72–82. 10.1107/s0907444905036693.

Evans, P.R., and Murshudov, G.N. (2013). How good are my data and what is the resolution? Acta Crystallogr D Biol Crystallogr 69, 1204–1214. 10.1107/s0907444913000061.

Feng, S., Knödler, A., Ren, J., Zhang, J., Zhang, X., Hong, Y., Huang, S., Peränen, J., and Guo, W. (2012). A Rab8 guanine nucleotide exchange factor-effector interaction network regulates primary ciliogenesis. The Journal of biological chemistry 287, 15602–15609. 10.1074/jbc.M111.333245.

Ferreira, R.R., Pakula, G., Klaeyle, L., Fukui, H., Vilfan, A., Supatto, W., and Vermot, J. (2018). Chiral Cilia Orientation in the Left-Right Organizer. Cell reports 25, 2008–2016.e2004. 10.1016/j.celrep.2018.10.069.

Fremont, S., Hammich, H., Bai, J., Wioland, H., Klinkert, K., Rocancourt, M., Kikuti, C., Stroebel, D., Romet-Lemonne, G., Pylypenko, O., et al. (2017). Oxidation of F-actin controls the terminal steps of cytokinesis. Nat Commun 8, 14528. 10.1038/ncomms14528.

Gautreau, A.M., Fregoso, F.E., Simanov, G., and Dominguez, R. (2022). Nucleation, stabilization, and disassembly of branched actin networks. Trends in cell biology 32, 421–432. 10.1016/j.tcb.2021.10.006.

Goetz, S.C., and Anderson, K.V. (2010). The primary cilium: a signalling centre during vertebrate development. Nat Rev Genet 11, 331–344. 10.1038/nrg2774.

Grimes, D.T., and Burdine, R.D. (2017). Left-Right Patterning: Breaking Symmetry to Asymmetric Morphogenesis. Trends Genet 33, 616–628. 10.1016/j.tig.2017.06.004.

Hashimoto, M., Shinohara, K., Wang, J., Ikeuchi, S., Yoshiba, S., Meno, C., Nonaka, S., Takada, S., Hatta, K., Wynshaw-Boris, A., and Hamada, H. (2010). Planar polarization of node cells determines the rotational axis of node cilia. Nature cell biology 12, 170–176. 10.1038/ncb2020.

Hauptmann, G., and Gerster, T. (1994). Two-color whole-mount in situ hybridization to vertebrate and Drosophila embryos. Trends Genet 10, 266. 10.1016/0168-9525(90)90008-t.

Hernandez-Hernandez, V., Pravincumar, P., Diaz-Font, A., May-Simera, H., Jenkins, D., Knight, M., and Beales, P.L. (2013). Bardet-Biedl syndrome proteins control the cilia length through regulation of actin polymerization. Hum Mol Genet 22, 3858–3868. 10.1093/hmg/ddt241.

Hildebrandt, F., Benzing, T., and Katsanis, N. (2011). Cil-iopathies. N Engl J Med 364, 1533–1543. 10.1056/NE-JMra1010172.

Hirokawa, N., Tanaka, Y., Okada, Y., and Takeda, S. (2006). Nodal flow and the generation of left-right asymmetry. Cell 125, 33–45. 10.1016/j.cell.2006.03.002.

Hoffman, H.K., and Prekeris, R. (2022). Roles of the actin cytoskeleton in ciliogenesis. J Cell Sci 135. 10.1242/jcs.259030.

Homma, Y., Hiragi, S., and Fukuda, M. (2021). Rab family of small GTPases: an updated view on their regulation and functions. Febs j 288, 36–55. 10.1111/febs.15453.

Jewett, C.E., Soh, A.W.J., Lin, C.H., Lu, Q., Lencer, E., Westlake, C.J., Pearson, C.G., and Prekeris, R. (2021). RAB19 Directs Cortical Remodeling and Membrane Growth for Primary Ciliogenesis. Developmental cell 56, 325–340.e328. 10.1016/j.devcel.2020.12.003.

Juhl, A.D., Anvarian, Z., Kuhns, S., Berges, J., Andersen, J.S., Wüstner, D., and Pedersen, L.B. (2023). Transient accumulation and bidirectional movement of KIF13B in primary cilia. J Cell Sci 136. 10.1242/jcs.259257.

Kabsch, W. (2010). XDS. Acta Crystallogr D Biol Crystallogr 66, 125–132. 10.1107/s0907444909047337.

Kim, J., Jo, H., Hong, H., Kim, M.H., Kim, J.M., Lee, J.K., Heo, W.D., and Kim, J. (2015). Actin remodelling factors control ciliogenesis by regulating YAP/TAZ activity and vesicle trafficking. Nat Commun 6, 6781. 10.1038/ncomms7781.

Kim, J., Lee, J.E., Heynen-Genel, S., Suyama, E., Ono, K., Lee, K., Ideker, T., Aza-Blanc, P., and Gleeson, J.G. (2010). Functional genomic screen for modulators of ciliogenesis and cilium length. Nature 464, 1048–1051. nature08895 [pii] 10.1038/nature08895.

Kimmel, C.B., Ballard, W.W., Kimmel, S.R., Ullmann, B., and Schilling, T.F. (1995). Stages of embryonic development of the zebrafish. Dev Dyn 203, 253–310. 10.1002/aja.1002030302.

Klena, N., and Pigino, G. (2022). Structural Biology of Cilia and Intraflagellar Transport. Annual review of cell and developmental biology 38, 103–123. 10.1146/annurev-cellbio-120219-034238.

Klinkert, K., and Echard, A. (2016). Rab35 GTPase: a central regulator of phosphoinositides and F-actin in endocytic recycling and beyond. Traffic. 10.1111/tra.12422.

Klinkert, K., Rocancourt, M., Houdusse, A., and Echard, A. (2016). Rab35 GTPase couples cell division with initiation of epithelial apico-basal polarity and lumen opening. Nat Commun 7, 11166. 10.1038/ncomms11166.

Knödler, A., Feng, S., Zhang, J., Zhang, X., Das, A., Peränen, J., and Guo, W. (2010). Coordination of Rab8 and Rab11 in primary ciliogenesis. Proceedings of the National Academy of Sciences of the United States of America 107, 6346–6351. 10.1073/pnas.1002401107.

Kobayashi, H., Etoh, K., and Fukuda, M. (2014a). Rab35 is translocated from Arf6-positive perinuclear recycling endosomes to neurite tips during neurite outgrowth. Small GT-Pases 5, e29290. 10.4161/sgtp.29290.

Kobayashi, H., Etoh, K., Ohbayashi, N., and Fukuda, M. (2014b). Rab35 promotes the recruitment of Rab8, Rab13 and Rab36 to recycling endosomes through MICAL-L1 during neurite outgrowth. Biol Open 3, 803–814. 10.1242/bio.20148771.

Kobayashi, H., and Fukuda, M. (2012). Rab35 regulates Arf6 activity through centaurin-beta2 (ACAP2) during neurite outgrowth. J Cell Sci 125, 2235–2243. 10.1242/jcs.098657.

Kobayashi, H., and Fukuda, M. (2013). Rab35 establishes the EHD1-association site by coordinating two distinct effectors during neurite outgrowth. J Cell Sci 126, 2424–2435. 10.1242/jcs.117846.

Konarev PV, Volkov V, Sokolova AV, Koch MHJ, and Di, S. (2003). PRIMUS: a Windows PC-based system for small-angle scattering data analysis. J. Appl. Cryst. 36, 1277–1282.

Kouranti, I., Sachse, M., Arouche, N., Goud, B., and Echard, A. (2006). Rab35 regulates an endocytic recycling pathway essential for the terminal steps of cytokinesis. Current biology: CB 16, 1719–1725.

Kuhns, S., Seixas, C., Pestana, S., Tavares, B., Nogueira, R., Jacinto, R., Ramalho, J.S., Simpson, J.C., Andersen, J.S., Echard, A., et al. (2019). Rab35 controls cilium length, function and membrane composition. EMBO Rep 20, e47625. 10.15252/embr.201847625.

Kuijl, C., Pilli, M., Alahari, S.K., Janssen, H., Khoo, P.S., Ervin, K.E., Calero, M., Jonnalagadda, S., Scheller, R.H., Neefjes, J., and Junutula, J.R. (2013). Rac and Rab GTPases dual effector Nischarin regulates vesicle maturation to facilitate survival of intracellular bacteria. The EMBO journal 32, 713–727. 10.1038/emboj.2013.10.

Kumar, R., Francis, V., Kulasekaran, G., Khan, M., Armstrong, G.A.B., and McPherson, P.S. (2022). A cell-based GEF assay reveals new substrates for DENN domains and a role for DENND2B in primary ciliogenesis. Sci Adv 8, eabk3088. 10.1126/sciadv.abk3088.

Kushawah, G., Hernandez-Huertas, L., Abugattas-Nuñez Del Prado, J., Martinez-Morales, J.R., DeVore, M.L., Hassan, H., Moreno-Sanchez, I., Tomas-Gallardo, L., Diaz-Moscoso, A., Monges, D.E., et al. (2020). CRISPR-Cas13d Induces Efficient mRNA Knockdown in Animal Embryos. Developmental cell 54, 805–817.e807. 10.1016/j.devcel.2020.07.013.

Legrand, P. (2017). XDSME: XDS Made Easier. GitHub repository https://github.com/legrandp/xdsme. 10.5281/zen-odo.837885.

Lu, H., Toh, M.T., Narasimhan, V., Thamilselvam, S.K., Choksi, S.P., and Roy, S. (2015a). A function for the Joubert syndrome protein Arl13b in ciliary membrane extension and ciliary length regulation. Dev Biol 397, 225–236. 10.1016/j.ydbio.2014.11.009.

Lu, Q., Insinna, C., Ott, C., Stauffer, J., Pintado, P.A., Rahajeng, J., Baxa, U., Walia, V., Cuenca, A., Hwang, Y.S., et al. (2015b). Early steps in primary cilium assembly require EHD1/EHD3-dependent ciliary vesicle formation. Nature cell biology 17, 228–240. 10.1038/ncb3109.

Madhivanan, K., and Aguilar, R.C. (2014). Cil-iopathies: the trafficking connection. Traffic 15, 1031–1056. 10.1111/tra.12195.

Magistrati, E., Maestrini, G., Niño, C.A., Lince-Faria, M., Beznoussenko, G., Mironov, A., Maspero, E., Bettencourt-Dias, M., and Polo, S. (2022). Myosin VI regulates ciliogenesis by promoting the turnover of the centrosomal/satellite protein OFD1. EMBO Rep 23, e54160. 10.15252/embr.202154160.

Malicki, J.J., and Johnson, C.A. (2017). The Cilium: Cellular Antenna and Central Processing Unit. Trends in cell biology 27, 126–140. 10.1016/j.tcb.2016.08.002.

Mitchison, H.M., and Valente, E.M. (2017). Motile and non-motile cilia in human pathology: from function to phenotypes. J Pathol 241, 294–309. 10.1002/path.4843.

Mizuno, K., Shiozawa, K., Katoh, T.A., Minegishi, K., Ide, T., Ikawa, Y., Nishimura, H., Takaoka, K., Itabashi, T., Iwane, A.H., et al. (2020). Role of Ca(2+) transients at the node of the mouse embryo in breaking of left-right symmetry. Sci Adv 6, eaba1195. 10.1126/sciadv.aba1195.

Molla-Herman, A., Ghossoub, R., Blisnick, T., Meunier, A., Serres, C., Silbermann, F., Emmerson, C., Romeo, K., Bourdoncle, P., Schmitt, A., et al. (2010). The ciliary pocket: an endocytic membrane domain at the base of primary and motile cilia. J Cell Sci 123, 1785–1795. 10.1242/jcs.059519.

Mul, W., Mitra, A., and Peterman, E.J.G. (2022). Mechanisms of Regulation in Intraflagellar Transport. Cells 11. 10.3390/cells11172737.

Nachury, M.V., Loktev, A.V., Zhang, Q., Westlake, C.J., Per-anen, J., Merdes, A., Slusarski, D.C., Scheller, R.H., Bazan, J.F., Sheffield, V.C., and Jackson, P.K. (2007). A core complex of BBS proteins cooperates with the GTPase Rab8 to promote ciliary membrane biogenesis. Cell 129, 1201–1213.

Nager, A.R., Goldstein, J.S., Herranz-Pérez, V., Portran, D., Ye, F., Garcia-Verdugo, J.M., and Nachury, M.V. (2017). An Actin Network Dispatches Ciliary GPCRs into Extracellular Vesicles to Modulate Signaling. Cell 168, 252–263.e214. 10.1016/j.cell.2016.11.036.

Nozaki, S., Katoh, Y., Terada, M., Michisaka, S., Funabashi, T., Takahashi, S., Kontani, K., and Nakayama, K. (2017). Regulation of ciliary retrograde protein trafficking by the Joubert syndrome proteins ARL13B and INPP5E. J Cell Sci 130, 563–576. 10.1242/jcs.197004.

Ojeda Naharros, I., and Nachury, M.V. (2022). Shedding of ciliary vesicles at a glance. J Cell Sci 135. 10.1242/jcs.246553.

Phua, S.C., Chiba, S., Suzuki, M., Su, E., Roberson, E.C., Pusapati, G.V., Schurmans, S., Setou, M., Rohatgi, R., Reiter, J.F., et al. (2017). Dynamic Remodeling of Membrane Composition Drives Cell Cycle through Primary Cilia Excision. Cell 168, 264–279.e215. 10.1016/j.cell.2016.12.032.

Phuyal, S., and Farhan, H. (2019). Multifaceted Rho GTPase Signaling at the Endomembranes. Front Cell Dev Biol 7, 127. 10.3389/fcell.2019.00127.

Pitaval, A., Tseng, Q., Bornens, M., and Théry, M. (2010). Cell shape and contractility regulate ciliogenesis in cell cycle-arrested cells. The Journal of cell biology 191, 303–312. 10.1083/jcb.201004003.

Rahajeng, J., Giridharan, S.S., Cai, B., Naslavsky, N., and Caplan, S. (2012). MICAL-L1 is a tubular endosomal membrane hub that connects Rab35 and Arf6 with Rab8a. Traffic 13, 82–93. 10.1111/j.1600-0854.2011.01294.x.

Read, R.J., and McCoy, A.J. (2011). Using SAD data in Phaser. Acta Crystallogr D Biol Crystallogr 67, 338–344. 10.1107/s0907444910051371.

Reiter, J.F., and Leroux, M.R. (2017). Genes and molecular pathways underpinning ciliopathies. Nature reviews. Molecular cell biology 18, 533–547. 10.1038/nrm.2017.60.

Robert, A., Margall-Ducos, G., Guidotti, J.E., Brégerie, O., Celati, C., Bréchot, C., and Desdouets, C. (2007). The intraflagellar transport component IFT88/polaris is a centrosomal protein regulating G1-S transition in non-ciliated cells. J Cell Sci 120, 628–637. 10.1242/jcs.03366.

Rodríguez, D., Sammito, M., Meindl, K., de Ilarduya, I.M., Potratz, M., Sheldrick, G.M., and Usón, I. (2012). Practical structure solution with ARCIMBOLDO. Acta Crystallogr D Biol Crystallogr 68, 336–343. 10.1107/s0907444911056071.

Rossi, A., Kontarakis, Z., Gerri, C., Nolte, H., Hölper, S., Krüger, M., and Stainier, D.Y. (2015). Genetic compensation induced by deleterious mutations but not gene knockdowns. Nature 524, 230–233. 10.1038/nature14580.

Sander, E.E., ten Klooster, J.P., van Delft, S., van der Kammen, R.A., and Collard, J.G. (1999). Rac downregulates Rho activity: reciprocal balance between both GT-Pases determines cellular morphology and migratory behavior. The Journal of cell biology 147, 1009–1022. 10.1083/jcb.147.5.1009.

Satir, P., Pedersen, L.B., and Christensen, S.T. (2010). The primary cilium at a glance. J Cell Sci 123, 499–503. 10.1242/jcs.050377.

Schneidman-Duhovny, D., Hammel, M., Tainer, J.A., and Sali, A. (2016). FoXS, FoXSDock and MultiFoXS: Single-state and multi-state structural modeling of proteins and their complexes based on SAXS profiles. Nucleic acids research 44, W424–429. 10.1093/nar/gkw389.

Schrödinger, L., and DeLano, W. (2020). PyMOL. Avail-able at: http://www.pymol.org/pymol.

Shaughnessy, R., and Echard, A. (2018). Rab35 GTPase and cancer: Linking membrane trafficking to tumorigenesis. Traffic. 10.1111/tra.12546.

Sheldrick, G.M. (2008). A short history of SHELX. Acta Crystallogr A 64, 112–122. 10.1107/s0108767307043930.

Simunovic, M., Voth, G.A., Callan-Jones, A., and Bassereau, P. (2015). When Physics Takes Over: BAR Proteins and Membrane Curvature. Trends in cell biology 25, 780–792. 10.1016/j.tcb.2015.09.005.

Smith, C.E.L., Lake, A.V.R., and Johnson, C.A. (2020). Primary Cilia, Ciliogenesis and the Actin Cytoskeleton: A Little Less Resorption, A Little More Actin Please. Front Cell Dev Biol 8, 622822. 10.3389/fcell.2020.622822.

Sorokin, S.P. (1968). Reconstructions of centriole formation and ciliogenesis in mammalian lungs. J Cell Sci 3, 207–230. 10.1242/jcs.3.2.207.

Stewart, K., Gaitan, Y., Shafer, M.E., Aoudjit, L., Hu, D., Sharma, R., Tremblay, M., Ishii, H., Marcotte, M., Stanga, D., et al. (2016). A Point Mutation in p190A RhoGAP Affects Ciliogenesis and Leads to Glomerulocystic Kidney Defects. PLoS genetics 12, e1005785. 10.1371/journal.pgen.1005785.

Streets, A.J., Prosseda, P.P., and Ong, A.C. (2020). Polycystin-1 regulates ARHGAP35-dependent centrosomal RhoA activation and ROCK signaling. JCI Insight 5. 10.1172/jci.insight.135385.

Svergun DI (1992). Determination of the regularization parameter in indirect-transform methods using perceptual criteria. J. Appl. Cryst. 25, 495–503.

Tate, J.G., Bamford, S., Jubb, H.C., Sondka, Z., Beare, D.M., Bindal, N., Boutselakis, H., Cole, C.G., Creatore, C., Dawson, E., et al. (2019). COSMIC: the Catalogue Of Somatic Mutations In Cancer. Nucleic acids research 47, D941–d947. 10.1093/nar/gky1015.

Varadi, M., Anyango, S., Deshpande, M., Nair, S., Natassia, C., Yordanova, G., Yuan, D., Stroe, O., Wood, G., Laydon, A., et al. (2022). AlphaFold Protein Structure Database: massively expanding the structural coverage of protein-sequence space with high-accuracy models. Nucleic acids research 50, D439–d444. 10.1093/nar/gkab1061.

Vonrhein, C., Flensburg, C., Keller, P., Sharff, A., Smart, O., Paciorek, W., Womack, T., and Bricogne, G. (2011). Data processing and analysis with the autoPROC toolbox. Acta Crystallogr D Biol Crystallogr 67, 293–302. 10.1107/s0907444911007773.

Wang, K., Yuen, S.T., Xu, J., Lee, S.P., Yan, H.H., Shi, S.T., Siu, H.C., Deng, S., Chu, K.M., Law, S., et al. (2014). Whole-genome sequencing and comprehensive molecular profiling identify new driver mutations in gastric cancer. Nat Genet 46, 573–582. 10.1038/ng.2983.

Webb, B., and Sali, A. (2016). Comparative Protein Structure Modeling Using MODELLER. Curr Protoc Bioinformatics 54, 5.6.1-5.6.37. 10.1002/cpbi.3.

Westlake, C.J., Baye, L.M., Nachury, M.V., Wright, K.J., Ervin, K.E., Phu, L., Chalouni, C., Beck, J.S., Kirkpatrick, D.S., Slusarski, D.C., et al. (2011). Primary cilia membrane assembly is initiated by Rab11 and transport protein particle II (TRAP-PII) complex-dependent trafficking of Rabin8 to the centrosome. Proceedings of the National Academy of Sciences of the United States of America 108, 2759–2764. 10.1073/pnas.1018823108.

Wheway, G., Lord, J., and Baralle, D. (2019). Splicing in the pathogenesis, diagnosis and treatment of ciliopathies. Biochim Biophys Acta Gene Regul Mech 1862, 194433. 10.1016/j.bbagrm.2019.194433.

Wheway, G., Nazlamova, L., and Hancock, J.T. (2018). Signaling through the Primary Cilium. Front Cell Dev Biol 6, 8. 10.3389/fcell.2018.00008.

Wheway, G., Schmidts, M., Mans, D.A., Szymanska, K., Nguyen, T.T., Racher, H., Phelps, I.G., Toedt, G., Kennedy, J., Wunderlich, K.A., et al. (2015). An siRNA-based functional genomics screen for the identification of regulators of ciliogenesis and ciliopathy genes. Nature cell biology 17, 1074–1087. 10.1038/ncb3201.

Wingfield, J.L., Lechtreck, K.F., and Lorentzen, E. (2018). Trafficking of ciliary membrane proteins by the intraflagellar transport/BBSome machinery. Essays Biochem 62, 753–763. 10.1042/ebc20180030.

Winn, M.D., Ballard, C.C., Cowtan, K.D., Dodson, E.J., Emsley, P., Evans, P.R., Keegan, R.M., Krissinel, E.B., Leslie, A.G., McCoy, A., et al. (2011). Overview of the CCP4 suite and current developments. Acta Crystallogr D Biol Crystallogr 67, 235–242. 10.1107/s0907444910045749.

Wu, C.T., Chen, H.Y., and Tang, T.K. (2018). Myosin-Va is required for preciliary vesicle transportation to the mother centriole during ciliogenesis. Nature cell biology 20, 175–185. 10.1038/s41556-017-0018-7.

Xie, S., Farmer, T., Naslavsky, N., and Caplan, S. (2019). MICAL-L1 coordinates ciliogenesis by recruiting EHD1 to the primary cilium. J Cell Sci 132. 10.1242/jcs.233973.

Yoder, B.K., Tousson, A., Millican, L., Wu, J.H., Bugg, C.E., Jr., Schafer, J.A., and Balkovetz, D.F. (2002). Polaris, a protein disrupted in orpk mutant mice, is required for assembly of renal cilium. Am J Physiol Renal Physiol 282, F541–552. 10.1152/ajprenal.00273.2001.

Youn, J.Y., Dunham, W.H., Hong, S.J., Knight, J.D.R., Bashkurov, M., Chen, G.I., Bagci, H., Rathod, B., MacLeod, G., Eng, S.W.M., et al. (2018). High-Density Proximity Mapping Reveals the Subcellular Organization of mRNA-Associated Granules and Bodies. Mol Cell 69, 517–532.e511. 10.1016/j.molcel.2017.12.020.

Yuan, S., Zhao, L., Brueckner, M., and Sun, Z. (2015). Intraciliary calcium oscillations initiate vertebrate left-right asymmetry. Current biology : CB 25, 556–567. 10.1016/j.cub.2014.12.051.

